# Rescue of Tomato spotted wilt tospovirus entirely from cDNA clones, establishment of the first reverse genetics system for a segmented (-)RNA plant virus

**DOI:** 10.1101/680900

**Authors:** Mingfeng Feng, Ruixiang Cheng, Minglong Chen, Rong Guo, Luyao Li, Zhike Feng, Jianyan Wu, Li Xie, Jian Hong, Zhongkai Zhang, Richard Kormelink, Xiaorong Tao

**Author notes:** Author contributions: M.F., Z.F. and X.T. conceived and designed the experiments and H.J., Z.Z. and R. K. provided input. M.F., R.C., M.C., G.R., L.L., Z.F., J.W., and X.L. performed the experiments. M.F., R. K. and X.T. wrote the manuscript.

## Abstract

The group of negative strand RNA viruses (NSVs) includes not only dangerous pathogens of medical importance but also serious plant pathogens of agronomical importance. Tomato spotted wilt tospovirus (TSWV) is one of those plant NSVs that cause severe diseases on agronomic crops and pose major threats to global food security. Its negative-strand segmented RNA genome has, however, always posed a major obstacle to molecular genetic manipulation. In this study, we report the complete recovery of infectious TSWV entirely from cDNA clones, the first reverse genetics (RG) system for a segmented plant NSV. First, a replication and transcription competent mini-genome replication system was established based on 35S-driven constructs of the S_(-)_-genomic (g) or S_(+)_-antigenomic (ag) RNA template, flanked by a 5’ Hammerhead and 3’ Ribozyme sequence of Hepatitis Delta virus, a nucleocapsid (N) protein gene and codon-optimized viral RNA dependent RNA polymerase (RdRp) gene. Next, a movement competent mini-genome replication system was developed based on M_(-)_-gRNA, which was able to complement cell-to-cell and systemic movement of reconstituted ribonucleoprotein complexes (RNPs) of S RNA replicon. After further optimization, infectious TSWV and derivatives carrying eGFP reporters were successfully rescued *in planta* via simultaneous expression of full-length cDNA constructs coding for S_(+)_-agRNA, M_(-)_-gRNA and L_(+)_-agRNA. Viral rescue occurred in the additional presence of various viral suppressors of RNAi, but TSWV NSs interfered with the rescue of genomic RNA. The establishment of a RG system for TSWV now allows detailed molecular genetic analysis of all aspects of tospovirus life cycle and their pathogenicity.

**Significance:** For many different animal-infecting segmented negative-strand viruses (NSVs), a reverse genetics system has been established that allows the generation of mutant viruses to study disease pathology and the role of *cis*- and *trans*-acting elements in the virus life cycle. In contrast to the relative ease to establish RG systems for animal-infecting NSVs, establishment of such system for the plant-infecting NSVs with a segmented RNA genome so far has not been successful. Here we report the first reverse genetics system for a segmented plant NSV, the Tomato spotted wilt tospovirus, a virus with a tripartite RNA genome. The establishment of this RG system now provides us with a new and powerful platform to study their disease pathology during a natural infection.

## Introduction

Negative-stranded RNA viruses (NSVs) present of a large group of viruses that include well known members of medical importance such as Ebola (EBOV), Rabies (RV), Influenza A (FLUAV) and Rift valley fever virus (RVFV) (1, 2). Infections with these viruses may cause considerable morbidity and mortality in humans and form an important burden on national health care budgets. The group also contains plant viruses of agronomical importance such as Tomato spotted wilt virus (TSWV) and Rice stripe virus (RSV) that cause severe diseases on agronomic crops and pose major threats to global food security (3–10).

Tospoviruses belong to the NSV with a segmented (tripartite) RNA genome and rank among the most devastating plant viruses worldwide (11, 12). They are classified in the family of *Tospoviridae* within the order *Bunyavidales* (13). TSWV is the type member of the only genus *Orthotospovirus*, in the family of *Tospoviridae* (11, 13). TSWV has very broad host range infecting more than one thousand plant species over 80 families (14) and is transmitted by thrips in a persistent, propagative manner (6, 9, 15, 16). Crops losses due to this virus have been estimated more than one billion dollars annually (7, 14).

TSWV consists of spherical, enveloped virus particles (80-120 nm) that contain a tripartite genome consisting of a large- (L), medium- (M), and small-sized (S) RNA segment (11). The L segment is of entire negative polarity, whereas the S and M segments are ambisense. The L segment encodes the viral RNA-dependent RNA polymerase (RdRp, ∼330 kDa) that is required for viral RNA replication and mRNA transcription (17, 18). The viral (v) strand of the M segment encodes the precursor to the glycoproteins (Gn and Gc, with n and c referring to the amino- and carboxy-terminal end of the precursor, respectively) in the negative sense and a nonstructural protein (NSm) in the positive sense. The glycoproteins are required for particle maturation and are presented as spikes on the surface of the virus envelope membrane (19, 20). They also play a major role as determinants for thrips vector transmission (21). The NSm plays pivotal roles in cell-to-cell and long distance movement of TSWV (22–26). The vRNA of the S segment codes for the nucleocapsid protein (N) in the negative sense and a nonstructural protein (NSs) in the positive sense. The N protein participates in the formation of ribonucleoprotein complexes (RNPs) (27–29) and is required for viral intracellular movement (30, 31). The NSs protein functions as a RNA silencing suppressor to defend against the plant innate immunity system (32–34), and triggers a defence response and concomittant programmed cell-death mediated by the dominant resistance gene *Tsw* from *Capsicum chinense* (35–37).

As a virus documented for almost a century (7, 38), TSWV has served as an important model for studying the molecular biology of tospovirus and other plant NSVs with segmented genomes (6–8, 11). However, its negative-strand tripartite RNA genome has posed a major obstacle to genetic manipulation of the virus. To initiate an infection cycle with this virus requires at least RNPs, the minimal infectious units that consist of viral RNA encapsidated by the N protein and associated with a few copies of the viral RNA-dependent RNA polymerase (6, 11). TSWV RNPs can be mechanically transferred from infected to healthy plants, however, transmission by thrips requires RNPs to be enveloped and spiked with the glycoproteins (21).

The first animal-infecting and related counterpart of TSWV with a segmented RNA genome to be rescued entirely from cDNA was from Bunyamwera virus in 1996 (39). Following this study, soon other segmented NSV were rescued from plasmid DNA. The influenza virus, containing a genome of eight RNA segments, was recovered in 1999 (40), while the first Arenavirus, with a bipartite RNA genome, was recovered in 2006 (41). Just recently, the first nonsegmented plant NSVs from the *Mononegavirales* have been rescued, *i.e*. the Sonchus yellow net nucleorhabdovirus (SYNV) (42, 43). A recent study also reported on the establishment of a TSWV S RNA-based mini-replicon in yeast (44), but in contrast to replication, no transcriptional activity was observed.

In contrast to the relative ease to establish RG systems for animal-infecting NSVs, reconstitution of infectious RNPs *in planta* for the plant-infecting viruses with a segmented RNA genome seems particularly difficult. Not only have all DNA constructs to be delivered into one and the same plant cell but for TSWV the RdRp is exceptionally large (∼330 kDa) compared to the RdRp of most other related bunyaviruses (∼240-260 kDa) and to typical open reading frames (ORFs) from the plant genome. Expression of such a large protein gene may not only be very inefficient but mRNA transcripts resulting from RNA polymerase II transcription of 35S promoter-constructs in the nucleus may also face splicing of cryptic splicing sites. Moreover, achieving proper ratios of all three genome segments in plant cells is not easily and consistently achieved by agrobacterium-mediated delivery of several constructs, which will affect the outcome of each individual experiment. All these obstacles may hamper the construction of a reverse genetics system for TSWV in plants.

In this study, we report the complete recovery of infectious TSWV entirely from cDNA clones in plants, the first reverse genetics system for a segmented plant NSV. The establishment of this system presents the start of a new research era for TSWV and provides us an entirely new and powerful platform to study the basic principles of the tospovirus life cycle and viral pathogenicity.

## Results

### Development of a TSWV S_(-)_-genomic RNA mini-replicon system in *Nicotiana benthamiana*

Prior to the rescuing of TSWV entirely from cDNA clones, a mini-replicon system based on the S RNA-template was established. To this end, a DNA copy of the TSWV S_(-)_-genomic RNA (gRNA) was cloned and flanked with a self-cleaving hammerhead (HH) ribozyme at the 5’-terminus and a hepatitis delta virus (HDV) ribozyme at the 3’-terminus. For visual monitoring, quantification purposes, and discrimination between primary and secondary genome transcription, the NSs and N genes were replaced with mCherry and eGFP, respectively (Fig. 1*A*). The resulting S_(-)_ mini-replicon reporter was cloned in a binary vector pCB301 downstream a double 35S promoter (2×35S) and denoted 35S:SR_(-)mCherry&eGFP_ (Fig. 1*A*), or a T7 promoter and denoted T7: SR_(-)mCherry&eGFP_ (Fig. S1*A*). The RdRp and N ORFs were amplified from cDNA of TSWV infected tissue and cloned into pCambia 2300 binary vector downstream a double 35S promoter. Binary vector constructs of the RdRp, N and four viral RNA silencing suppressor genes (VSRs: NSs from TSWV, P19 from Tomato bushy stunt tombusvirus (TBSV), HcPro from Tobacco etch potyvirus (TEV) and γb from Barley yellow mosaic hordeivirus (BYMV)) were agroinfiltrated in *N. benthamiana* either with 35S:SR_(-)mCherry&eGFP_ or with T7:SR_(-)mCherry&eGFP_ and a 35S-driven T7 RNA polymerase gene construct and next monitored for eGFP fluorescence. However, during repeated experiments no eGFP fluorescence was observed from 35S:SR_(-)mCherry&eGFP_ nor T7:SR_(-)mCherry&eGFP_ (Fig. S1*B* and *C*).

**Fig. 1.**
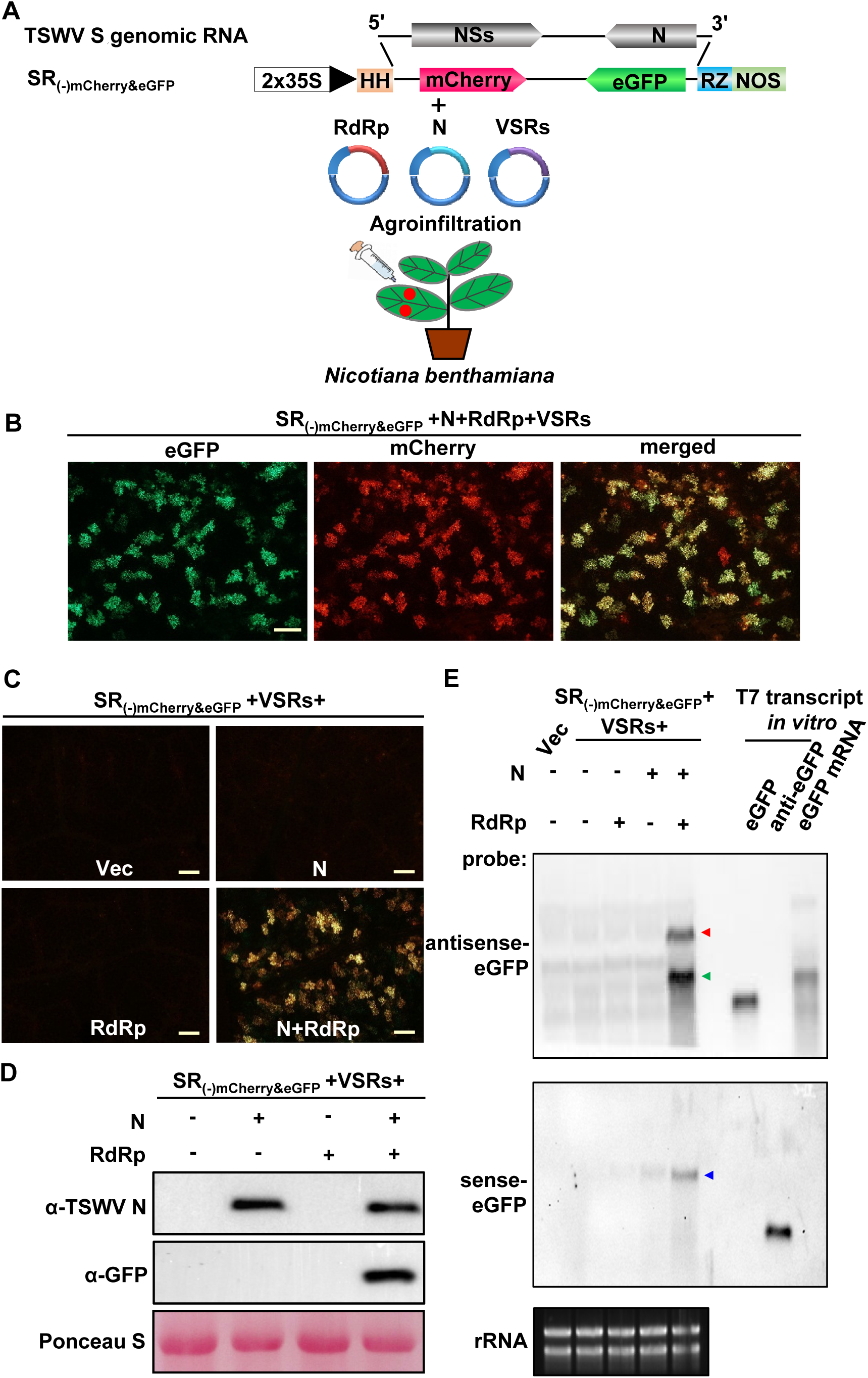
Construction of a TSWV S_(-)_ RNA-based mini-replicon system in *N. benthamiana*. (*A*) Schematic representation of binary constructs to express TSWV S_(-)_ mini-replicon, TSWV N, RdRp and four RNA silencing suppressors (VSRs: NSs, P19, HcPro and γb) proteins by agroinfiltration into *N. benthamiana*. The S_(-)_-gRNA of TSWV is shown on the top. SR_(-)mCherry&eGFP_: the NSs and N of S_(-)_-gRNA were replaced by mCherry and eGFP, respectively. (-) refers to the negative (genomic)-strand of S RNA; 2×35S: a double 35S promoter; HH: hammerhead ribozyme; RZ: Hepatitis Delta virus (HDV) ribozyme; NOS: nopaline synthase terminator. (*B*) Foci of eGFP and mCherry fluorescence in *N. benthamiana* leaves co-expressing SR_(-)mCherry&eGFP_, RdRp, N and four VSRs at 5 days post infiltration (dpi) under a fluorescence microscope. The bar represents 400 μm. (*C*) Analysis of RdRp and N requirement for SR_(-)mCherry&eGFP_ mini-genome replication in *N. benthamiana* leaves. SR_(-)mCherry&eGFP_ was coexpressed with pCB301 empty vector (Vec), N, RdRp or both in *N. benthamiana* leaves by agroinfiltration. Agro-infiltrated leaves were examined and photographed at 5 dpi under a fluorescence microscope. Signal shown reflects a merge of mCherry and eGFP fluorescence from both reporter genes. Bar represents 400 μm. (*D*) Immunoblot analysis on the expression of N and eGFP proteins in the leaves shown in panel (*C*) using specific antibodies against N and GFP, respectively. Ponceau S staining of rubisco large subunit is shown for protein loading control. (*E*) Northern blot analysis of S_(-)_-mini-replicon replication and transcription in the presence of N, RdRp or both in *N. benthamiana*. The S RNA genomic, anti-genomic and subgenomic transcripts (eGFP mRNA) were detected by DIG-labeled sense eGFP or anti-sense eGFP probes. The red and blue arrows indicate the anti-genomic and genomic RNAs of SR_(-)mCherry&eGFP_, respectively. The green arrow indicates the eGFP mRNA transcript. Ethidium bromide staining of ribosomal RNA (rRNA) was used as RNA loading control.

The possibility of failures to establish a mini-replicon system for TSWV could be due to low (unstable) expression of the TSWV RdRp protein, therefore, the codon usage of the RdRp gene was optimized for *N. benthamiana* and potential intron splicing sites were removed. The optimized RdRp gene (RdRp_opt_) was cloned in a binary, 35S-driven expression vector and next, again agroinfiltrated in *N. benthamiana* leaves together with binary expression constructs of the N gene, the four VSR gene constructs and the 35S:SR_(-)mCherry&eGFP_ mini-replicon reporter. At 5 days post infiltration (dpi), expression of the reporter genes was analyzed by monitoring for mCherry and eGFP fluorescence in the *N. benthamiana* leaves (Fig. 1*A* and *B*). Whereas no fluorescence was observed in the controls, *i.e.* leaves agroinfiltrated with 35S:SR_(-)mCherry&eGFP_ alone or co-expressing 35S:SR_(-)mCherry&eGFP_ with RdRp_opt_ or N only, eGFP and mCherry fluorescence was consistently observed in leaves agroinfiltrated with 35S:SR_(-)mCherry&eGFP_ and both N and RdRp_opt_ (Fig. 1*C*). This was confirmed by Western immunoblot analysis (Fig. 1*D*).

Northern blot analysis showed that both the genomic RNA and anti-genomic RNA of the SR_(-)mCherry&eGFP_ mini-replicon were detected in the leaves that co-expressed both N and RdRp (here after RdRp represent optimized RdRp) at 5 dpi, but not in the leaves co-expressing RdRp or N only (Fig. 1*E*). In addition, using an anti-sense eGFP probe genome length S RNA subgenomic-sized RNA, likely presenting eGFP transcripts, were detected (Fig. 1*E*, upper panel). A time course analysis showed that eGFP and mCherry fluorescence in *N. benthamiana* leaves was visible from 3 dpi onwards and gradually increased to 12 dpi (Fig. S2*A*). This was also confirmed by immunoblot analysis (Fig. S2*B*).

Altogether the results indicated that in *N. benthamiana* the 35S replicon transcript SR_(-)mCherry&eGFP_ was properly processed by the HH and RZ, and next used by the RdRp as template for primary transcription of mRNA_eGFP_, as well replicated into SR_(-)mCherry&eGFP_ for secondary transcription of mRNA_mCherry_. Furthermore, the codon-optimized RdRp clearly exhibited full functionality in supporting viral genome transcription and replication, while the wild type RdRp for unknown reasons didn’t. When the T7:SR_(-)mCherry&eGFP_ mini-replicon reporter was co-expressed with T7 RNA polymerase, RdRp, N and four VSRs in *N. benthamiana* leaves, somewhat unexpected, no mCherry or eGFP fluorescence was detected (Fig. S1*C* and *D*).

### The optimization of the concentration of N and RdRp, and VSRs on TSWV SR_(-)mCherry&eGFP_ mini-replicon

Having established a TSWV S_(-)_-gRNA based mini-replicon system in *N. benthamiana*, attempts were made to further optimize the system. To this end, binary constructs of the TSWV S_(-)_ mini-replicon were agroinfiltrated into *N. benthamiana* in the additional presence of varying amounts of N and RdRp gene expression constructs. During these experiments the amounts of *Agrobacterium* carrying the N expression construct were increased (from OD_600_ 0.2 to 0.8) while the *Agrobacterium* harboring the RdRp expression construct was kept fixed at OD_600_ 0.2, and vice versa. The results showed highest eGFP reporter gene expression from the 35S:SR_(-)mCherry&eGFP_ mini-replicon when *Agrobacterium* suspensions bearing N and RdRp expression constructs were both infiltrated at OD_600_ 0.2. When *Agrobacterium* harboring either N or RdRp was infiltrated onto *N. benthamiana* leaves at OD_600_ >0.4 the expression of eGFP from the S_(-)_-mini-replicon greatly decreased (Fig. 2 *A*, *B* and *C*). Furthermore, at high concentrations (OD_600_ > 0.6) of *Agrobacterium*, the visible cell death was triggered in the infiltrated leaves (data not shown).

**Fig. 2.**
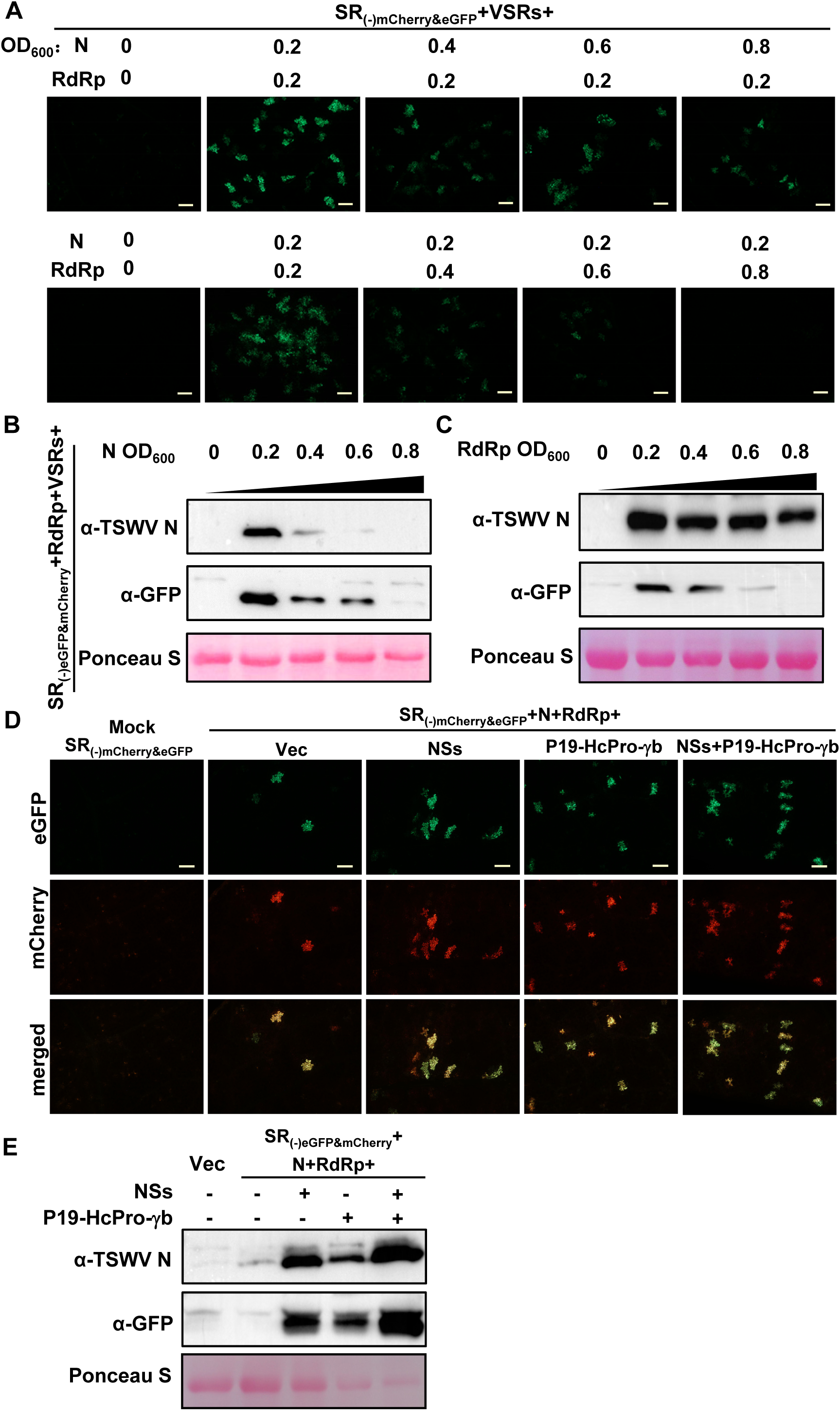
Optimization of the SR_(-)mCherry&eGFP_ mini-replicon system. (*A*) Optimizing the concentration of N and RdRp proteins for replication and transcription of SR_(-)mCherry&eGFP_ in *N. benthamiana* leaves. Increasing amounts of *Agrobacterium,* from OD_600_= 0.2 to 0.8 and containing the binary expression constructs for N (upper panels) or RdRp (bottom panels), were mixed with fixed amounts of *Agrobacterium* containing the RdRp or N construct (OD_600_ 0.2), respectively, and their effects on eGFP reporter expression were visualized under a fluorescence microscope at 5 dpi. Bars represent 400 μm. (*B*) and (*C*) Western immunoblot detection of the N and eGFP proteins expressed in the leaves shown in panel (*A*) using specific antibodies against N and GFP, respectively. (*D*) Optimization of RNA silencing suppressors (VSRs) on SR_(-)mCherry&eGFP_ mini-reporter replication and transcriptions as measured by eGFP and mCherry expression. The SR_(-)mCherry&eGFP_, N and RdRp proteins were co-expressed with pCB301 empty vector (Vec), NSs, P19-HcPro-γb or all four VSRs in *N. benthamiana* leaves. Foci expressing eGFP and mCherry in agroinfiltrated leaves were visualized under a fluorescence microscope at 5 dpi. Bars represent 400 μm. (*E*) Western immunoblot detection of N and eGFP protein synthesis in the leaves shown in panel (*D*) using N and GFP-specific antibodies, respectively. Ponceau S staining was used as protein loading control.

Next, in a similar approach and using the optimized setting, the effects of VSRs on the replication and transcription of the SR_(-)mCherry&eGFP_ mini-replicon were investigated. Without the addition of VSRs, mCherry and eGFP fluorescence was only observed in a small number of cells, but these numbers increased in the addition of TSWV NSs and/or the three VSRs P19, HcPro and γb. The largest number of cells showing eGFP expression from the S_(-)_ mini-replicon, as monitored by fluorescence, were obtained when all four VSRs were added (Fig. 2*D* and *E*). These observations were further confirmed by immunoblot assays (Fig. 2*E*).

### The role of *cis*-acting sequences in transcription and replication of the TSWV SR_(-)mCherry&eGFP_ replicon

Using the optimized SR_(-)mCherry&eGFP_ mini-replicon system, the role of the 5’- untranslated region (5’-UTR), 3’-UTR and the non-coding A/U-rich intergenic region (IGR) between NSs and N genes (45, 46) in replication-transcription was examined. To this end, SR_(-)mCherry&eGFP_ derivatives were made from which the 5’-UTR, IGR or 3’-UTR, respectively, were deleted and next tested on transcription-replication using the mini-replicon assay (Fig. S3*A*). No eGFP and mCherry fluorescence was observed when the 5’-UTR or 3’-UTR of SR_(-)mCherry&eGFP_ was removed. However, eGFP reporter expression could still be detected when the IGR of SR_(-)mCherry&eGFP_ was deleted (Fig. S3*B*). Immunoblot analysis confirmed the expression of eGFP from SR_(-)mCherry&eGFPΔIGR_ (ΔIGR), and lack of expression from SR_(-)mCherry&eGFPΔ5’UTR_ (Δ5’UTR) and SR_(-)mCherry&eGFPΔ3’UTR_ (Δ3’UTR) (Fig. S3*C*). To analyze whether the lack of eGFP expression was a matter of translation or transcription-replication, samples from infiltrated *N. benthamiana* were collected and analyzed by Northern blot. The results showed that for SR_(-)mCherry&eGFPΔ5’UTR_ (Δ5’UTR) and SR_(-)mCherry&eGFPΔ3’UTR_ (Δ3’UTR), weak signals of genomic RNA could be detected but they are not similar to the signal strength obtained for the genome length and mRNA molecules as seen with the SR_(-)mCherry&eGFP_ replicon, while anti-genomic RNAs could not be detected (Fig. S3*D*). For SR_(-)mCherry&eGFPΔIGR_ (ΔIGR) both RNA strands were detected (Fig. S3*D*), suggesting that IGR is not essential for viral RNA synthesis, while no signal is obtained for the mRNA length molecules as seen with the SR_(-)mCherry&eGFP_ replicon. Taken together, these findings suggest that the 5’-UTR and 3’-UTR play an essential role in viral transcription and replication of TSWV RNA segments.

### Development of a TSWV S_(+)_-agRNA based mini-replicon system

For many reverse genetics systems of NSV, mini-replicons have initially been established based on gRNA (vRNA). Here, we managed to develop a first system for TSWV based on antigenomic (ag)RNA (vcRNA). In order to analyze whether a system could be developed based on agRNA, a S_(+)_-agRNA mini-replicon was constructed similarly to the one based on S_(-)_-gRNA but in which the N gene was maintained and only the NSs gene was replaced by eGFP, denoted SR_(+)eGFP_ (Fig. 3*A*). In analogy to the replicon assays with SR_(-)mCherry&eGFP_, *N. benthamiana* leaves were agro-infiltrated with binary expression constructs of SR_(+)eGFP_, four VSRs and either N or RdRp separately or together, respectively, and monitored for eGFP fluorescence. Whereas eGFP fluorescence, resulting from primary transcription of the replicon transcript by viral RdRp, was not detected when SR_(+)eGFP_ was expressed alone or in the additional presence of N, eGFP fluorescence was observed when SR_(+)eGFP_ was co-expressed with both RdRp and N, but also when SR_(+)eGFP_ was co-expressed with RdRp alone (Fig. 3*B*). This strongly indicated that a certain (residual) amount of SR_(+)eGFP_ transcripts, resulting from 35S transcription, did not become fully processed by the HH and RZ and remained functional in translation, thereby giving rise to N protein. This was confirmed by Western immunoblot analysis (Fig. 3*C*). Northern blot analysis furthermore showed that samples from the replicon assays performed in the presence of RdRp and N or RdRp alone, besides eGFP mRNA transcripts, also contained agRNA and gRNA of SR_(+)eGFP_, indicating the occurrence of replication (Fig. 3*D*). Altogether, these results demonstrate that the N protein can also be expressed from the SR_(+)eGFP_ replicon to support its transcription and replication. This provides an attractive alternative to the S_(-)_-gRNA based mini-replicon as additional binary expression constructs for N do not have to be supplied anymore.

**Fig. 3.**
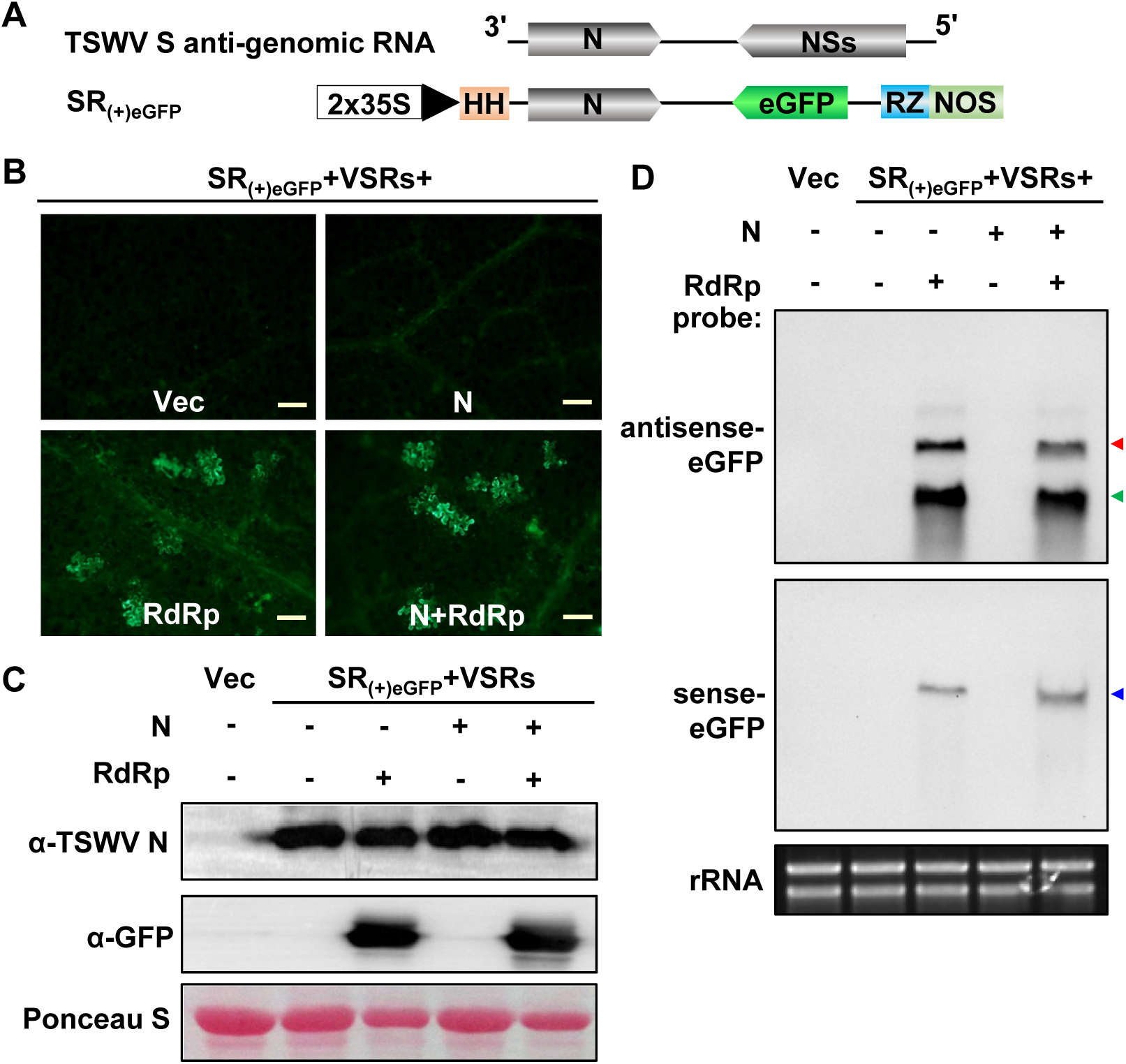
Development of the S_(+)_-agRNA mini-replicon system in *N. benthamiana*. (*A*) Schematic representation of the TSWV SR_(+)eGFP_ mini-replicon. The NSs gene of TSWV S agRNA was replaced by eGFP. Anti-genomic RNA strands of the SR_(+)eGFP_ mini-replicon are transcribed from a double 35S promoter (2×35S), and flanked by a HH ribozyme and HDV ribozyme (RZ) sequence. (+) refers to the positive (antigenomic)-strand of S RNA. (*B*) Foci of eGFP fluorescence in *N. benthamiana* leaves co-expressing TSWV SR_(+)eGFP_ with pCB301 empty vector (Vec), N, RdRp or N+RdRp by agroinfiltration. The agroinfiltrated leaves were photographed at 3 dpi under a fluorescence microscope. Bars represent 400 μm. (*C*) Western immunoblot detection of N and eGFP protein synthesis in the leaves shown in panel (*B*) using N- and GFP-specific antibodies, respectively. Ponceau S staining was used as a protein loading control. (D) Northern blot analysis of the replication and transcription of SR_(+)eGFP_ mini-replicon in *N. benthamiana* co-expressed with empty vector (Vec), N, RdRp or both. The anti-genomic RNAs (red arrow), genomic RNAs (blue arrow) and eGFP mRNA transcripts (green arrow) were detected with DIG-labeled sense and anti-sense eGFP probes, respectively. Ethidium bromide staining was used as RNA loading control.

### Development of a M_(-)_-gRNA based mini-replicon for cell-to-cell movement of TSWV in *N. benthamiana*

As a first step towards development of a reverse genetics system to rescue TSWV virus entirely from cDNA, a movement competent mini-replicon was also established. To this end, a TSWV M**_(-)_**-gRNA based mini-replicon was constructed, similar as the ones made for S_(-)_ and S_(+)_.Within this construct the NSm cell-to-cell movement protein gene was maintained but the GP ORF was exchanged for eGFP, resulting in a mini-replicon designated as MR_(-)eGFP_ (Fig. 4*A*). After *Agrobacterium*-mediated delivery into *N. benthamiana*, no eGFP fluorescence was observed in leaves containing the MR_(-)eGFP_ replicon with RdRp or N. However, in the presence of both RdRp and N, eGFP fluorescence was observed in many cells that connected each other (Fig. 4*B* and *C*). In comparison to the eGFP fluorescence always expressed in a single plant cells from SR_(-)mCherry&eGFP_ or SR_(+)eGFP_ reporter, the results suggested that the MR_(-)eGFP_ mini-replicon has moved from cell-to-cell in *N. benthamiana* leaves. Northern blot analysis confirmed the synthesis of gRNA, agRNA and (subgenomic-length) eGFP mRNA transcripts of the MR_(-)eGFP_ replicon in the presence of both RdRp and N, but not with RdRp or N only (Fig. 4*D*).

**Fig. 4.**
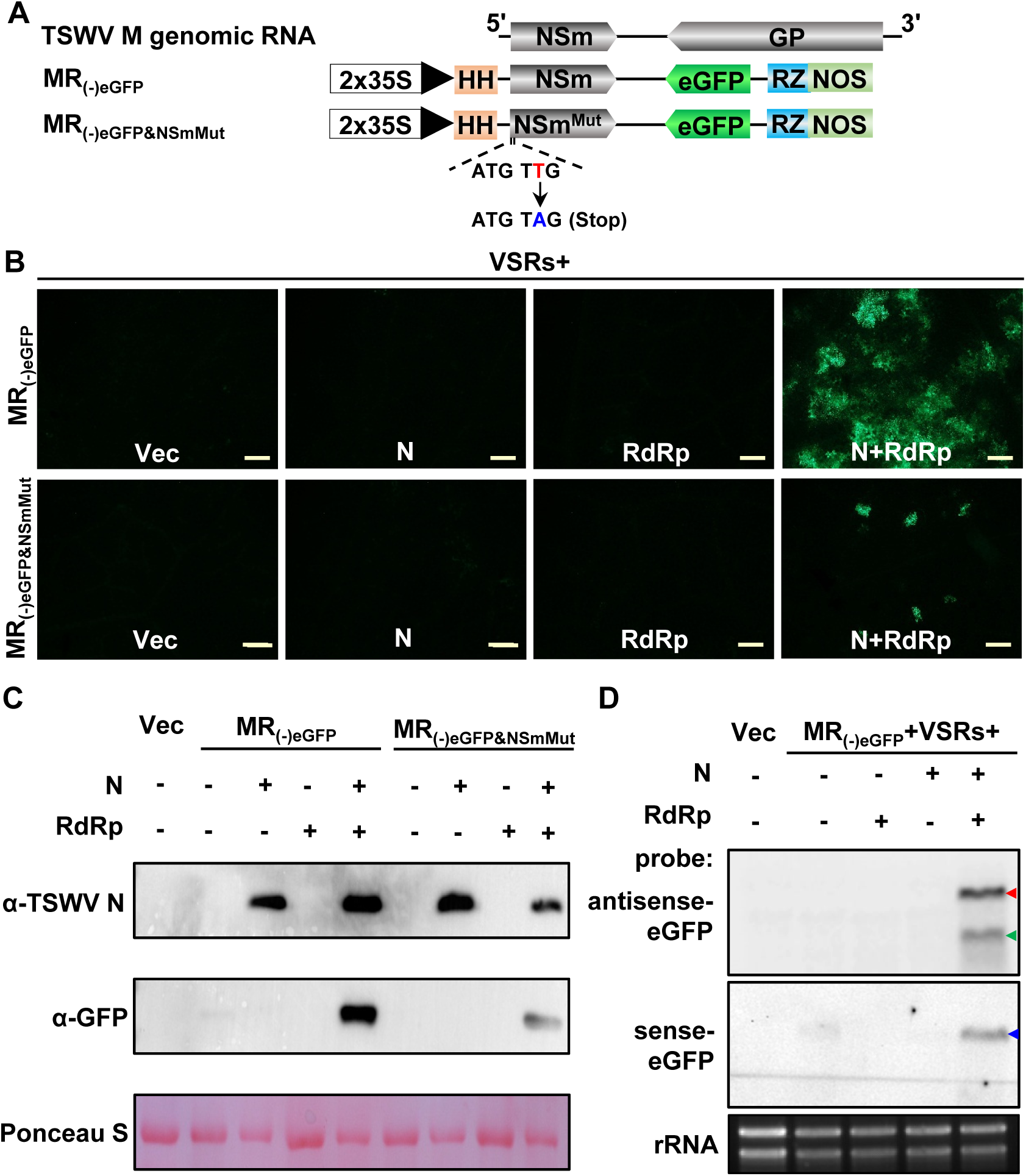
Establishment of a TSWV M_(-)_-gRNA based mini-replicon system with cell-to-cell movement competency in *N. benthamiana*. (*A*) Schematic representation of the TSWV MR_(-)eGFP_ mini-replicon and its mutant derivative MR_(-)eGFP&NSmMut_. The GP gene of TSWV M_(-)_ gRNA was substituted by eGFP. The genomic RNA of the mini-replicon is transcribed from a double 35S promoter (2×35S) and flanked by a Hammerhead (HH) ribozyme and HDV ribozyme (RZ). For MR_(-)eGFP&NSmMut_, a stop codon was introduced immediately after the start codon of the NSm gene in the MR_(-)eGFP_ mini-replicon. (*B*) Foci of eGFP fluorescence expressed from the MR_(-)eGFP_ or MR_(-)eGFP&NSmMut_ mini-replicon in *N. benthamiana* leaves co-expressed with the empty vector (Vec), N, RdRp or N+RdRp by agroinfiltration. Agroinfiltrated leaves were photographed at 4 dpi under a fluorescence microscope. Bars represent 400 μm. (*C*) Western immunoblot detection of N and eGFP protein synthesis in the leaves shown in panel (*B*) with specific antibodies against N and GFP, respectively. Ponceau S staining was used as protein loading control. (*D*) Northern blot analysis of the replication and transcription of the MR_(-)eGFP_ mini-replicon co-expressed with Vec, N, RdRp or N+RdRp in *N. benthamiana* leaves. The anti-genomic RNAs (red arrow), genomic RNAs (blue arrow) and eGFP mRNA transcripts (green arrow) were detected with DIG labeled sense and anti-sense eGFP probes, respectively. Ethidium bromide staining was used as RNA loading control.

To further substantiate the findings on possible cell-to-cell movement of the MR_(-)eGFP_ mini-replicon, a stop codon was introduced immediately downstream the start codon of NSm and the construct designated MR_(-)eGFP&NSmMut_) (Fig. 4*A*). When the MR_(-)eGFP&NSmMut_ replicon was delivered and co-expressed with RdRp and N in *N. benthamiana* leaves, eGFP fluorescence was only detected in a single cells (Fig. 4*B*). As expected, Western immunoblot analysis confirmed the production of eGFP protein in leaves containing the MR_(-)eGFP&NSmMut_ replicon, and in significantly lower amounts compared to the MR_(-)eGFP_ replicon (Fig. 4*C*).

### Establishment of the systemic infection of M_(-)_- and S_(+)_-mini-replicon reporters by co-expression of full-length antigenomic L_(+)_ in *N. benthamiana*

With the establishment of S (g/ag)RNA-based mini-replicon systems, and a movement-competent M gRNA-based mini-replicon, we set out to construct full length genomic cDNA clones of L_(-)_, M_(-)_ and S_(-)_, flanked by HH and HDV at 5’- and 3’-terminus, as a first step towards the rescue of TSWV entirely from cDNA clones. At the same time, similar constructs were made for the anti-genomic L_(+)_, M_(+)_ and S_(+)_. However, attempts to recover infectious TSWV from *N. benthamiana* after agrobacterium-mediated delivery of all binary expression constructs of L_(-)_, M_(-)_, S_(-)_ together with N, RdRp and four VSRs, but also with the anti-genomic L_(+)_, M_(+)_ and S_(+)_ constructs, all failed (Table 1).

**Table 1.**
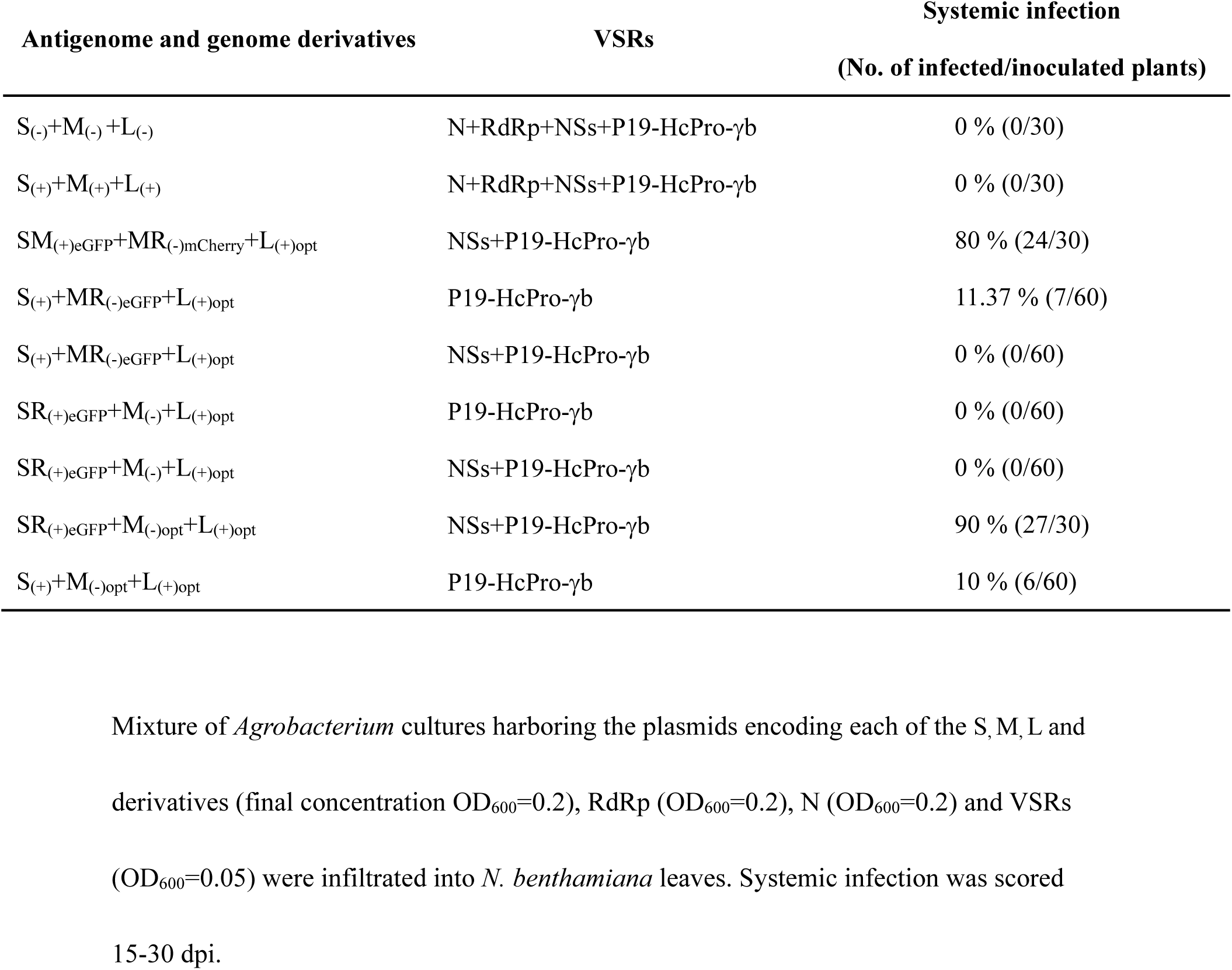
Systemic infection rate of recombinant TSWV rescued in N. benthamiana in the presence of viral suppressors of RNA silencing (VSRs).

Since MR_(-)eGFP_ was earlier shown to be movement competent, it was next investigated whether the M_(-)_- and S_(-)_-minigenomes moved into the same plant cell in the presence of both RdRp and N. Upon co-expression of MR_(-)mCherry_, SR_(-)eGFP_, RdRp and N in *N. benthamiana* leaves, expression of both mCherry and eGFP from the MR_(-)mCherry_ and SR_(-)eGFP_ mini-replicons, respectively, could be discerned. However, the foci of mCherry fluorescence were separate from those showing eGFP fluorescence (Fig. S4). Previously, ectopic expression of Tobacco crinkle virus (TCV) RdRp was reported to cause superinfection exclusion, and prevented the entry of progeny virus into the original cell expressing the RdRp (47). Ectopic expression of TSWV RdRp and N would possibly cause superinfection exclusions and block intercellular movement of SR_(-)eGFP_ into cells containing MR_(-)mCherry_, To avoid that, a new strategy was employed in which RdRp and N were expressed from viral agRNAs. To this end, a construct was made of the full-length L agRNA containing the optimized RdRp and flanked by the HH and HDV ribozymes, denoted L_(+)opt_ (Fig. 5*A*). To test the expression and functionality of RdRp from this construct, L_(+)opt_ was co-expressed with the SR_(-)mCherry&eGFP_ mini-replicon, N and VSRs in *N. benthamiana* leaves. The results showed clear eGFP and mCherry fluorescence and indicated that L_(+)opt_ was able to support SR_(-)mCherry&eGFP_ transcription and replication (Fig. S5*A*). Furthermore, L_(+)opt_ was also able to support the replication and transcription of the SR_(+)eGFP_ mini-replicon, without the additional ectopic expression of N (Fig. S5*B*), and of the movement competent MR_(-)eGFP_ (Fig. S5*C*).

**Fig. 5.**
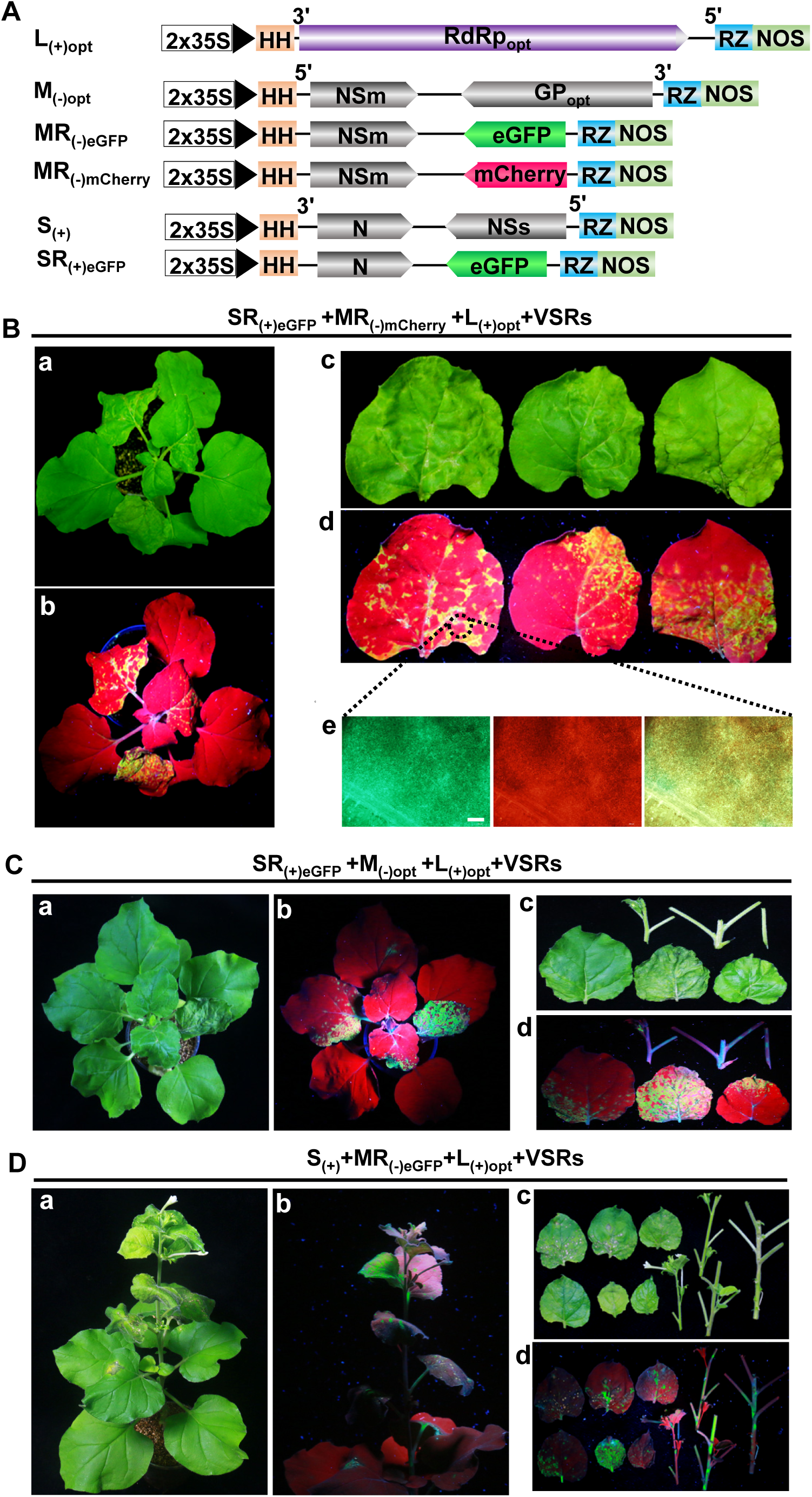
Establishment of a systemic infection in *N. benthamiana* with replicons S_(+)_ and M_(-)_ co-expressed with full length antigenomic L_(+)opt_. (*A*) Schematic representation of constructs expressing TSWV full length antigenomic L_(+)opt_ with optimized RdRp, full length genomic M_(-)opt_ with optimized GP, MR_(-)eGFP_, MR_(-)mCherry_, full length antigenomic S_(+)_ and SR_(+)eGFP_. Primary viral RNA transcripts are transcribed from a double 35S promoter (2×35S) and flanked by a HH and HDV ribozyme (RZ). (*B*) eGFP and mCherry fluorescence in *N. benthamiana* resulting from systemic infection of agroinfiltrated SR_(+)eGFP_, MR_(-)mCherry_ and L_(+)opt_ constructs. The systemic infected plants (*a* and *b*) and leaves (*c* and *d*) were photographed at 21 dpi under white light and (hand-held) ultraviolet (UV) light. Foci of eGFP and mCherry fluorescence in leaves shown in panel *d* as visualized under a fluorescence microscope. Bar represents 400 μm. (*C*) eGFP fluorescence in *N. benthamiana* resulting from systemic infection of agroinfiltrated SR_(+)eGFP_, M_(-)opt_ and L_(+)opt_ constructs. Infected plants (*a* and *b*) and leaves (*c* and *d*) were photographed at 18 dpi under white light and (hand-held) UV light, respectively. (*D*) eGFP fluorescence in *N. benthamiana* resulting from systemic infection of agroinfiltrated S_(+)_, MR_(-)eGFP_ and L_(+)opt_ constructs. Infected plants (*a* and *b*) and leaves (*c* and *d*) were photographed at 50 dpi under white light and (hand-held) UV light, respectively.

In a next experiment L_(+)opt_ was co-expressed with MR_(-)mCherry_, SR_(+)eGFP_ and four VSRs in *N. benthamiana* and plants analyzed for a systemic infection (Fig. 5*A*). At 6 dpi, mCherry and eGFP fluorescence were detected in the locally agroinfiltrated *N. benthamiana* leaves and in which some foci were found to express mCherry and eGFP together (Fig. S6*A*). At 15 dpi, necrotic symptoms became visual in systemic leaves of *N. benthamiana* (Fig. 5*B, a* and *c*). Using a handheld UV lamp, a clear eGFP fluorescence was also observed in those leaves (Fig. 5*B, b* and *d*). The eGFP signal was detected in 24 out of 30 agro-infiltrated *N. benthamiana* plants (Table 1). Both eGFP and mCherry fluorescence were detected in veins/stems and systemic leaves under a fluorescence microscope (Fig. 5*B*, *e*). The systemic infection of *N. benthamiana* with SR_(+)eGFP_, MR_(-)mCherry_ and L_(+)opt_ was further confirmed by RT-PCR analysis (Fig. S6*B*).

### Recovery of infectious TSWV from the full-length cDNA clones

Based on the establishment of a systemic infection after *Agrobacterium*-mediated delivery of replicons SR_(+)eGFP_, MR_(-)mCherry_ and L_(+)opt_, we next generated a full-length construct for S agRNA without any reporter gene, designated as S_(+)_ and co-expressed it with replicon constructs L_(+)opt_, M_(-)_ and four VSRs in *N. benthamiana* leaves. However, and surprisingly, no infectious TSWV was recovered from systemic leaves of *N. benthamiana* that were infiltrated with these constructs (Table 1). To find out whether this was due to failure of S_(+)_, we next examined whether S_(+)_ was able to establish a systemic infection in combination with the functional MR_(-)eGFP_ and L_(+)opt_ constructs. When L_(+)opt_, MR_(-)eGFP_, S_(+)_ and three VSRs (P19-HcPro-γb) were co-expressed in *N. benthamiana* leaves, eGFP fluorescence was visible at 18 dpi in systemic leaves of *N. benthamiana*, although not as efficient as in the case with the SR_(+)eGFP_ replicon, since only 7 out of 60 plants showed systemic infection (Fig. 5*D* and Table 1). RT-PCR analysis confirmed the systemic infection with S_(+)_, MR_(-)eGFP_ and L_(+)opt_ in those *N. benthamiana* (Fig. S6*C*). When L_(+)opt_, MR_(-)eGFP_ and S_(+)_ were co-expressed with four VSRs (P19-HcPro-γb and NSs) in *N. benthamiana* leaves, intriguingly, no eGFP fluorescence was observed in the systemic leaves (Table 1) suggesting that ectopic expression of NSs interfered with the rescue of virus from full length S_(+)_, MR_(-)eGFP_ and L_(+)opt_.

Next, we tested the rescuing of virus from full length M_(-)_, SR_(+)eGFP_ and L_(+)opt_. To this end, the constructs of L_(+)opt_, M_(-)_ and SR_(+)eGFP_ were delivered and co-expressed in *N. benthamiana* in the presence of either four (P19-HcPro-γb+NSs) or three (P19-HcPro-γb) VSRs. The results showed no eGFP fluorescence in systemic leaves of *N. benthamiana* at 15-50 dpi, indicating that M_(-)_ was not able to complement and rescue the S_(+)_-mini-replicons into systemic leaves (Table 1). Northern blot analysis showed that neither gRNAs nor agRNAs were detected for M_(-)_ while, in contrast, gRNAs and agRNAs were detected for S_(+)_ (Fig. S7*A* and *B*). Earlier, the MR_(-)eGFP_ mini-replicon was shown to replicate and transcribe (Fig. 4*D*). The only difference between M_(-)_ and MR_(-)eGFP_ mini-replicon was the GP gene, which was exchanged for eGFP in the second construct. Considering that primary M_(-)_ transcripts were produced in the nucleus by the 35S promoter, and putative splice sites were also predicted in the GP sequence (*SI Appendix,* Table S2), it was likely that primary M_(-)_ transcripts were prone to splicing before sufficient replication of the mini-replicon and transcriptional-translational expression of the cell-to-cell movement protein gene could take place. For this reason, codon optimization was performed on the GP gene sequence in M_(-)_, leading to a new construct designated as M_(-)opt_ (Fig. 5*A*). Upon co-expression of L_(+)opt_, M_(-)opt_ and the SR_(+)eGFP_ mini-replicon in *N. benthmiana* eGFP fluorescence was observed in systemic leaves (Fig. 5*C*). Fluorescence was observed in 27 out of 30 agroinfiltrated plants, and demonstrated that M_(-)opt_ produced a functional and stable M genomic RNA, able to replicate and support systemic movement of S and L RNP molecules by its encoded NSm protein (Table 1). RT-PCR analysis further confirmed a systemic infection of *N. benthamiana* with SR_(+)eGFP_, M_(-)opt_ and L_(+)opt_ (Fig. S6*D*).

In a final experiment, aiming to rescue “wild type” TSWV entirely from cDNA clones, the binary constructs of L_(+)opt_, M_(-)opt_ and S_(+)_ were agroinfiltrated into *N. benthamiana* leaves together with three VSRs (P19-HcPro-γb). At 19 dpi, typical leaf curling was observed in systemic leaves from *N. benthamiana* plants (Fig. 6*A* and Table 1). Upon disease progression, plants started to exhibit a stunted phenotype between 19-30 dpi. When the experiment was repeated with a large batch of plants, a systemic infection was observed in 6 out of 60 plants (Table 1). Northern blot analyses on samples collected from systemically infected leaves showed the presence of gRNA and agRNA of S, M and L RNA segments (Fig. 6*B*), which was also confirmed by RT-PCR (Fig. S8). Moreover, sequence analysis of the amplicons derived from the L and M RNA confirmed the presence of codon optimized RdRp and GP gene sequences (Fig. 6*C*). Immunoblot analysis on systemically infected leaf samples showed the presence of N, NSs, NSm, Gn and Gc proteins *N. benthamiana* (Fig. 6*D*), altogether indicating a successful systemic infection with rescued TSWV (rTSWV).

**Fig. 6.**
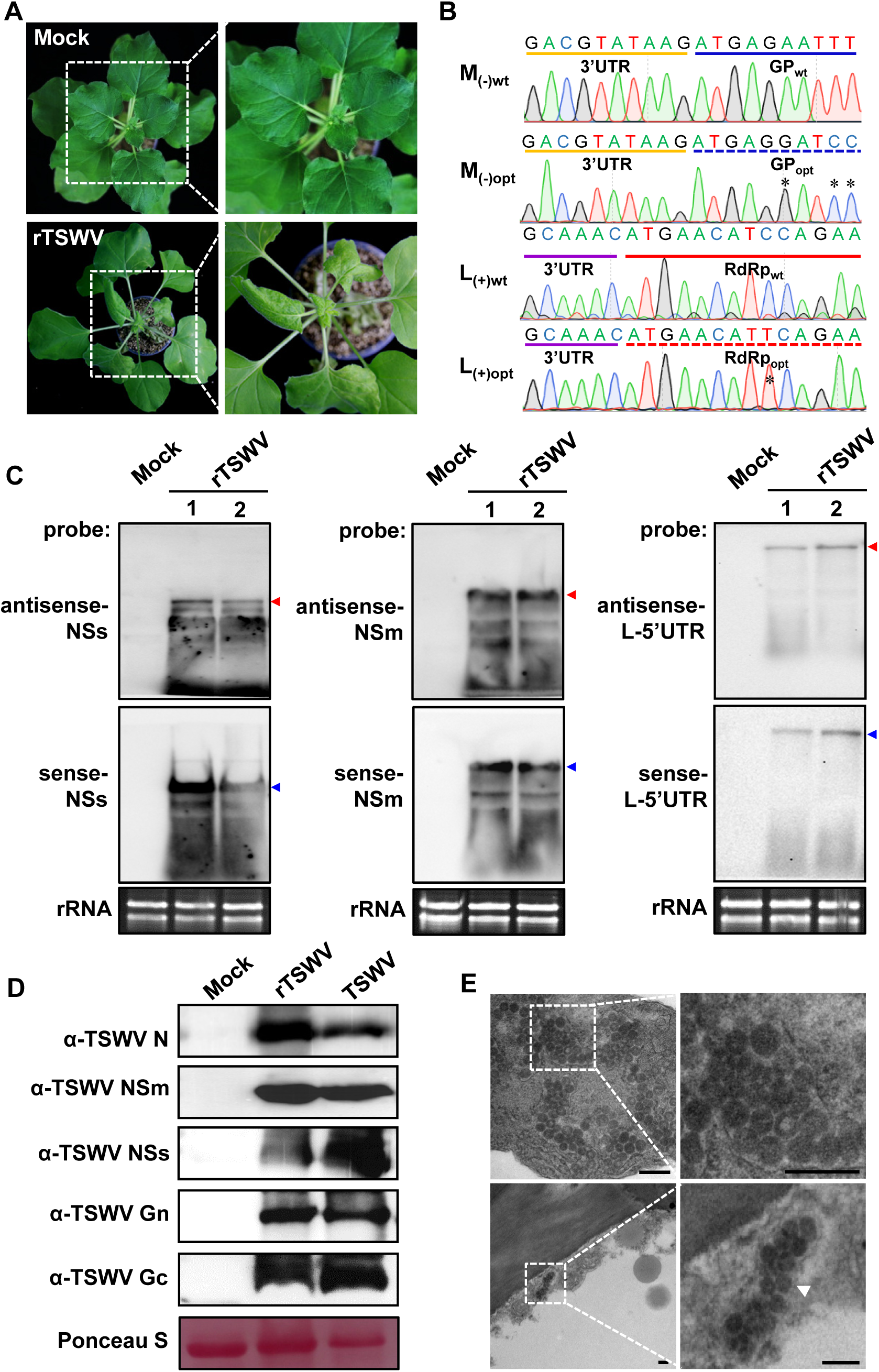
Rescue of infectious TSWV from full-length cDNA clones in *N. benthamiana*. (*A*) Systemic infection of *N. benthamiana* plants with rescued TSWV (rTSWV) resulting from agoinfiltration of S_(+)_, M_(-)opt_ and L_(+)opt_ and three VSRs (P19, HcPro and γb). The plant agroinfiltrated with pCB301 empty vector was used as a mock control. Images were taken at 19 dpi. Boxed areas of the plants that show stunting, mosaic and leaf curling are shown enlarged in the right panels. (*B*) Sequence confirmation of codon optimized sequences of the GP gene (from the M_(-)opt_ RNA segment) and the RdRp gene (from the L_(+)opt_ RNA segment) on RT-PCR fragments obtained from systemic leaves of *N. benthamiana* infected with rTSWV. The optimized sequence of GP from rTSWV is underlined by a blue dashed line, and wild type GP sequence underlined in blue. The 3’-untranslated region (UTR) sequence of the M genomic RNA is marked with a yellow line. The optimized sequence of RdRp from rTSWV is underlined by a red dashed line, and wild type RdRp sequence underlined in red. The 3’-UTR sequence of the L genomic RNA is marked with a purple line. The stars indicate the codon optimization sites of GP and RdRp gene sequences. (*C*) Northern blot detection of viral RNA from the S, M and L RNA segment, respectively, in systemic leaves of *N. benthamiana* infected with rTSWV. The anti-genomic RNAs (red arrow) and genomic RNAs (blue arrow) were detected with DIG labeled sense and anti-sense NSs-, NSm- and L-5’UTR probes, respectively. Lane 1 and 2 refer to two independent replicates. Ethidium bromide staining was used as RNA loading control. (*D*) Western immunoblot detection of the N, NSm, NSs, Gc, and Gn proteins from leaves systemically infected with rTSWV, using specific antibodies against N, NSm, NSs, Gc, and Gn, respectively. Leaves infected with wild-type TSWV were used as a positive control. Ponceau S staining was used as protein loading control. (*E*) Electron micrographs of thin sections of *N. benthamiana* plants infected with rTSWV. Boxed regions in the left panels and showing the presence of virions, are shown enlarged in the right panels. Spherical enveloped virus particles are indicated (white arrow head). Bars represent 0.2 μm.

To demonstrate genuine virus particle rescue of rTSWV, samples were collected from newly infected systemic leaf tissues and subjected to transmission electron microscopy. As shown in Fig. 6*E*, typical enveloped and spherical virus particles were observed in rTSWV-infected tissue, altogether indicating that infectious TSWV (rTSWV) was successfully rescued from full-length cDNA clones of L_(+)opt_, M_(-)opt_ and S_(+)_.

## Discussion

The establishment of a reverse genetics system for a segmented NSV basically requires two steps. The first one involves the *in vivo* reconstitution of transcriptionally active RNPs, often managed by development of a mini-genome replication system. The second step involves virus rescue entirely from full-length “infectious” cDNA clones, based on tools developed and optimized with the mini-genome replication system. In this study, we first successfully reconstituted infectious RNPs based on TSWV S_(-)_-gRNA and S_(+)_-agRNA after having optimized the sequence of RdRp. Next, a movement competent mini-genome replication system was developed based on M_(-)_-gRNA, which was also able to complement and systemically rescue reconstituted S RNPs. In a third step, full length constructs were made for S_(+)eGFP_-agRNA, M_(-)mCherry_-gRNA and L_(+)_-agRNA, to directly accommodate for translation of (small amounts of) all three genomic (35S) transcripts into N, NSm and RdRp proteins, respectively, and leave out the additional need of ectopically expressed N and RdRp. *Agrobacterium*-mediated delivery of these constructs lead to a successful systemic infection of *N. benthamiana* with rTSWV carrying eGFP reporters. In a last step, the GP gene sequence of M_(-)_ was optimized, that allowed the final rescuing of infectious rTSWV particles entirely from full-length cDNA clones in *N. benthamiana*.

The genomic RNAs of segmented NSVs possess neither a 5’ cap-structure nor 3’-poly(A) tail (2, 48). Instead, their termini contain highly conserved sequences that show inverted sequence complementary and fold into a panhandle structure with a major role in RNA transcription and replication. Any additional nucleotide residues at those termini in the past have been shown to disrupt/affect transcription-replication of animal-infecting segmented NSVs (49). For this reason, the choice of plant promoter to generate the first primary full-length genomic RNA templates (mimicking authentic genomic RNA molecules) for initiating viral replication is one of the major and critical factors for the construction of a reverse genetics system for TSWV. For animal-infecting segmented NSVs, researchers in the past have been using various systems. One of the first strategies employed bacteriophage T7 promoter constructs co-expressed with a T7 RNA polymerase and later followed by the use RNA polymerase I (Pol I) promoter constructs to generate the initial viral genome length RNA transcripts in mammalian cells (39, 40, 50–52). Unfortunately, attempts to establish the TSWV mini-replicon system based on the T7 promoter and T7 RNA polymerase strategy was unsuccessful (Fig. S1*A, C* and *D*). The activity of the Pol I promoter was shown to be species-dependent (53). Although a Pol I promoter has been reported from *Arabidopsis* (54, 55), while the transcription initiation +1 site is still not known. For *N. benthamiana* no Pol I promoter has been characterized yet. The 35S promoter, an RNA Pol II promoter, is well characterized and hence remains the only choice to establish a reverse genetics system for TSWV in plants. This is be in contrast to reverse genetics of segmented NSVs in animal cells, where all viruses have been reconstituted after T7/Pol I driven production of primary viral RNA templates for replication. The Pol II promoter has been used to produce the initial viral RNA transcripts of an animal-infecting nonsegmented NSV (56). Recently, the 35S/Pol II promoter was also successfully employed to produce primary viral RNA template of the first non-segmented plant NSV reconstituted, the SYNV rhabdovirus (42, 43). Here, we successfully deployed the 35S/Pol II promoter and two ribozymes at 5’ and 3’ ends of viral RNA sequences, to generate full length viral RNA transcripts that are recognized as initial/’’authentic’’ RNA templates for TSWV replication and transcription by viral N and RdRp.

Besides the right promoter, the RdRp protein may present another bottleneck for the establishment of a reverse genetics system. Tospoviruses code for a single, unprocessed ∼330 kDa RdRp from the 8.9 kb-sized L RNA (17, 18). The RdRp gene sequence of TSWV was predicted to contain numerous intron splicing sites (*SI Appendix,* Table S1). Since the first animal segmented negative strand RNA virus was rescued in 1996 (39), numerous groups worldwide have attempted to construct a reverse genetics system for a tospovirus in plants. Here, it is shown that codon optimization and removal of potential intron splicing sites have been crucial for the expression of a functional RdRp of tospovirus from 35S-driven constructs *in planta* (Fig. 1*B*). While codon optimization may have contributed to increased protein expression levels, removal of predicted potential intron splicing sites from the RdRp genemay have helped to further stabilize and increase expression levels. After all, TSWV is known to replicate in the cytoplasm (2, 48), and its RdRp gene may not have been evolved to escape from the nuclear (pre-mRNA) splicing machinery. However, after nuclear transcription of the RdRp gene by the 35S promoter, any intron splicing site in the wild type RdRp transcript could thus be spliced and result in a truncated, non-functional RdRp.

Not only for RdRp, but also an optimized GP gene sequence turned out to be crucial to rescue a full length M RNA-based transcriptionally active RNP. Whereas the M_(-)mCherry_ mini-replicon was able to establish a systemic infection in *N. benthamiana* when co-expressed with S_(+)eGFP_ mini-replicon and L_(+)opt_, the wild type full length M segment did not. Like in the case with the RdRp gene sequence, the GP gene sequence of TSWV was also predicted to contain numerous intron splicing sites (*SI Appendix,* Table S2). The absence of antigenomic and genomic RNA strands from the wild type full length M replicon on Northern blots (Fig. S7*B*) indicated the possibility that primary transcripts could have been prone to splicing in the GP sequence. This would not only lead to a loss of genome length RNA molecules, but also inhibit the production of NSm protein (either from direct translation of the primary M transcript, or after secondary transcription of NSm mRNA), needed for cell-to-cell and systemic movement of viral RNPs.

Not only the wild type sequence of L and M RNA segments may be spliced in the nucleus, also the S RNA segment generated by the 35S promoter could be prone to splicing. This is supported by the experiments in which *N. benthamiana* where infiltrated with the S_(+)eGFP_ mini-replicon, M_(-)opt_ and L_(+)opt_ and resulted in 80% virus recovery (Table 1), but when only the S_(+)eGFP_ mini-replicon was exchanged for the full length S_(+)_ virus recovery dropped to 11.37 %. The very same reason may explain the low infection rate (10 %) when full length S_(+)_, M_(-)opt_ and L_(+)opt_ are co-expressed (Table 1). Although this could be due to the splicing of S RNA, the (residual) levels of full length S produced apparently have been sufficient to initiate viral replication.

Similar to other bunyaviruses (39, 52, 57), both the RdRp and N proteins are required for reconstitution of infectious RNPs complexes for TSWV (Fig 1*C*). However, high expression of either N or RdRp results in cell death and cause negative effects on the replication and transcription of TSWV. Moreover, ectopic expression of N and RdRp by the strong 35S promoter also seems to cause superinfection exclusion, as earlier observed and reported with various viruses infecting humans, animals, and plants (47, 58–60). During superinfection exclusion a preexisting infection of virus prevents a secondary infection with the same or a highly similar virus. It is an active virus-controlled process that may be determined by a specific viral protein. For example, for potyvirus the coat protein and NIa protease have been identified to control superinfection exclusion (60). For TCV, the p28, involved in replication protein, was shown to confer superinfection exclusion as *a priori* expression of p28 blocked (re-)infection with TCV (47). Ectopic expression of N and RdRp may also have blocked progeny L-, M- and S- RNAs from moving into neighboring plant cells. However, the presence of L-, M- and S- RNA segments in the same cells is a pre-requisite for the reconstitution of infectious TSWV and to systemic spread in *N. benthamiana.* Fortunately, direct expression of N from S_(+)_ and RdRp from L_(+)opt_ have helped to overcome superinfection exclusion and to recover infectious TSWV. Whether this is due to the fact that N and RdRp are directly expressed from primary viral genome transcripts and simultaneously associate to progeny L-, M- and S- RNA segments in the same plant cells into infectious RNPs and/or involves a more fine-tuned protein expression relative to RNA segment replication remains unclear. Ectopic expression of NSs also inhibited the rescue of full length S_(+)_ segment from cDNA. Since TSWV NSs significantly enhanced the replication of S and M mini-replicons lacking the NSs ORF, this indicated that the inhibition could relate to (simultaneous / *a priori*) ectopic expression of NSs gene sequences with overlap to the full length S_(+)_ replicon. This could be tested by ectopic expression of an untranslatable NSs^ΔATG^ construct.

In summary, a series of issues has hampered the construction of a successful TSWV reverse genetics system: the choice of promoter and construct design to generate primary viral RNA transcripts in plants that mimick authentic viral RNA molecules, the expression of a very large viral RdRp, negative effects of ectopic expression of RdRp, N and NSs, and the absence of viral RNA synthesis of the wild type M_(-)_ segment. In this study, we have been able to solve all these issues and successfully managed to establish a reverse genetics system for the tripartite RNA genome of TSWV. Using the S RNA mini-replicon system containing eGFP and mCherry reporter genes, the role of *cis*-and *trans*-acting elements for viral replication and transcription can be studied. Using the M-RNA mini-replicon system cell-to-cell movement of TSWV RNPs *in planta* can be studied. To track the virus during systemic infection of plants rTSWV can be generated containing fluorescent reporter genes at the genetic loci of either GP or NSs. The establishment of this RG system now provides us with a new and powerful platform to generate mutant viruses and study their disease pathology in a natural setting, including basic principles of all tospovirus life cycle and viral pathogenicity. As a personal communication, Jeanmarie Verchot’s group has also recovered the Rose rosette emaravirus entirely from cDNA clones, a plant NSV with 7 RNA segments. The establishment of these RG systems presents the start of a new research era for the segmented plant NSVs.

## Materials and Methods

Details of the methodology used are provided in *SI Appendix,* Materials and Methods, and include plasmid construction, plant material and growth conditions, agro-infiltration, immunoblot analysis, Northern blot analysis, RT-PCR, GFP imaging, fluorescence microscopy and Electron microscopy. Primers used in this study are listed in *SI Appendix,* Table S3.

## ACKNOWLEDGMENTS

We thank Dr. Yi Xu for critical review of this manuscript. This work was supported by the National Natural Science Foundation of China (31630062 and 31471746), the Fundamental Research Funds for the Central Universities (JCQY201904), Youth Science and Technology Innovation Program to XT and Postgraduate Research & Practice Innovation Program of Jiangsu Province to MF.

## Supplemental Figure Legends

**Fig. S1.**
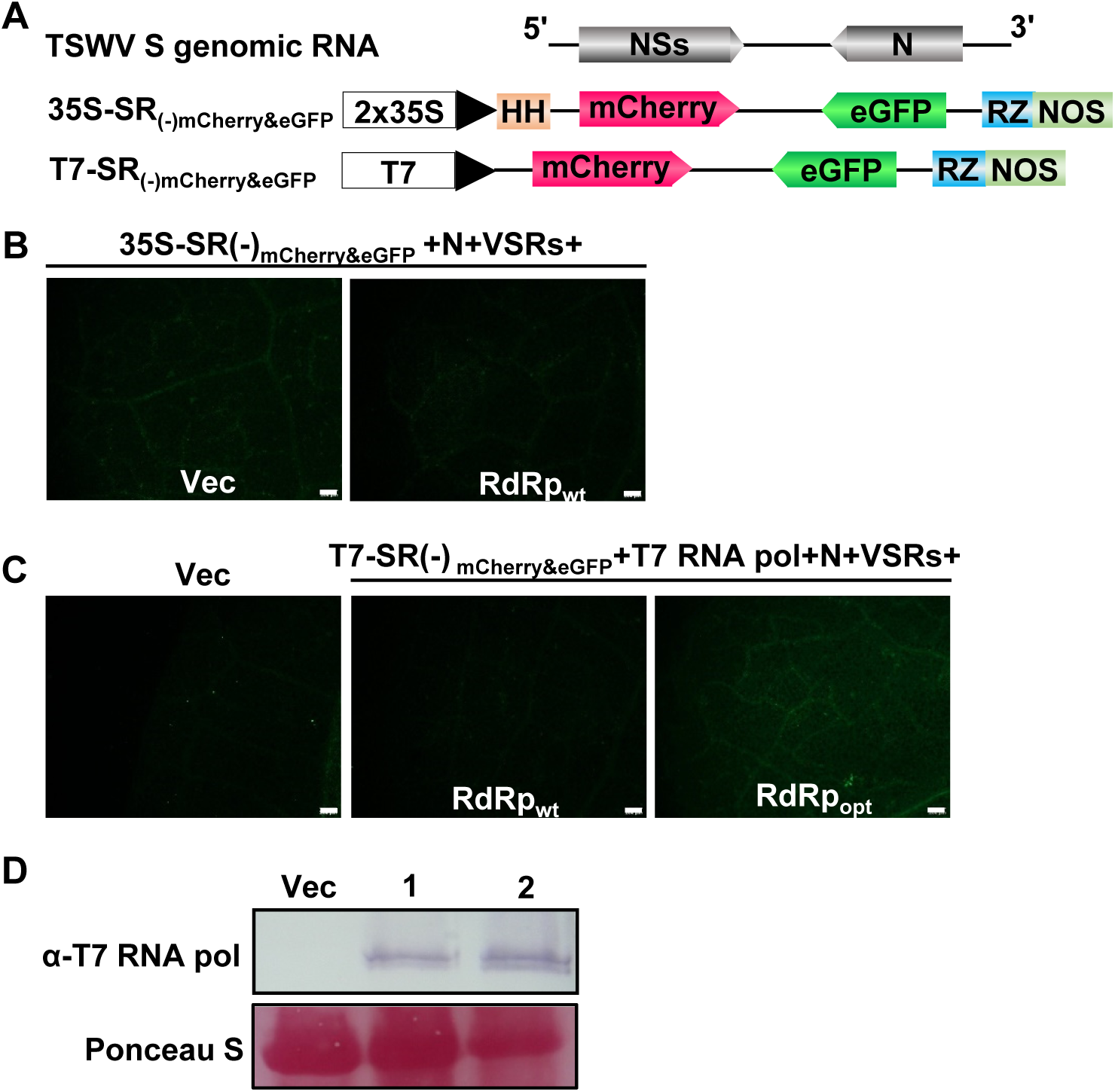
Functional analysis of wild type RdRp and the use of T7 promoter in a mini-genome replication assay. (*A*) Schematic diagram of TSWV 35S:SR_(-)mCherry&eGFP_ and T7:SR_(-)mCherry&eGFP_ mini-replicon reporters. (*B*) The wild type RdRp (RdRp_wt_) or the empty vector (Vec) was co-expressed with 35S:SR_(-)mCherry&eGFP_, N, VSRs (VSRs: NSs, P19, HcPro and γb) in *N. benthamiana* leaves. The expression of eGFP was examined by a fluorescence microscope. (*C*) Constructs coding for T7:SR_(-)mCherry&eGFP_, T7 RNA polymerase (pol), N and VSRs were co-expressed with RdRp_wt_ or RdRp_opt_ in *N. benthamiana* leaves. Replication of T7:SR_(-)mCherry&eGFP_ was examined by monitoring for eGFP fluorescence with a fluorescence microscope. Empty vector (Vec) pCB301 was used as a negative control. Bars represent 200 μm. (*D*) Western immunoblot detection of T7 RNA pol using a T7 RNA pol specific antibody. Ponceau S staining was used as protein loading control. Lane 1: sample from leaves co-expressing T7:SR_(-)mCherry&eGFP_, T7 RNA Pol, N, VSRs and RdRp_wt_; lane 2: sample from leaves co-expressing T7:SR_(-)mCherry&eGFP_, T7 RNA Pol, N, VSRs and RdRp_opt_.

**Fig. S2.**
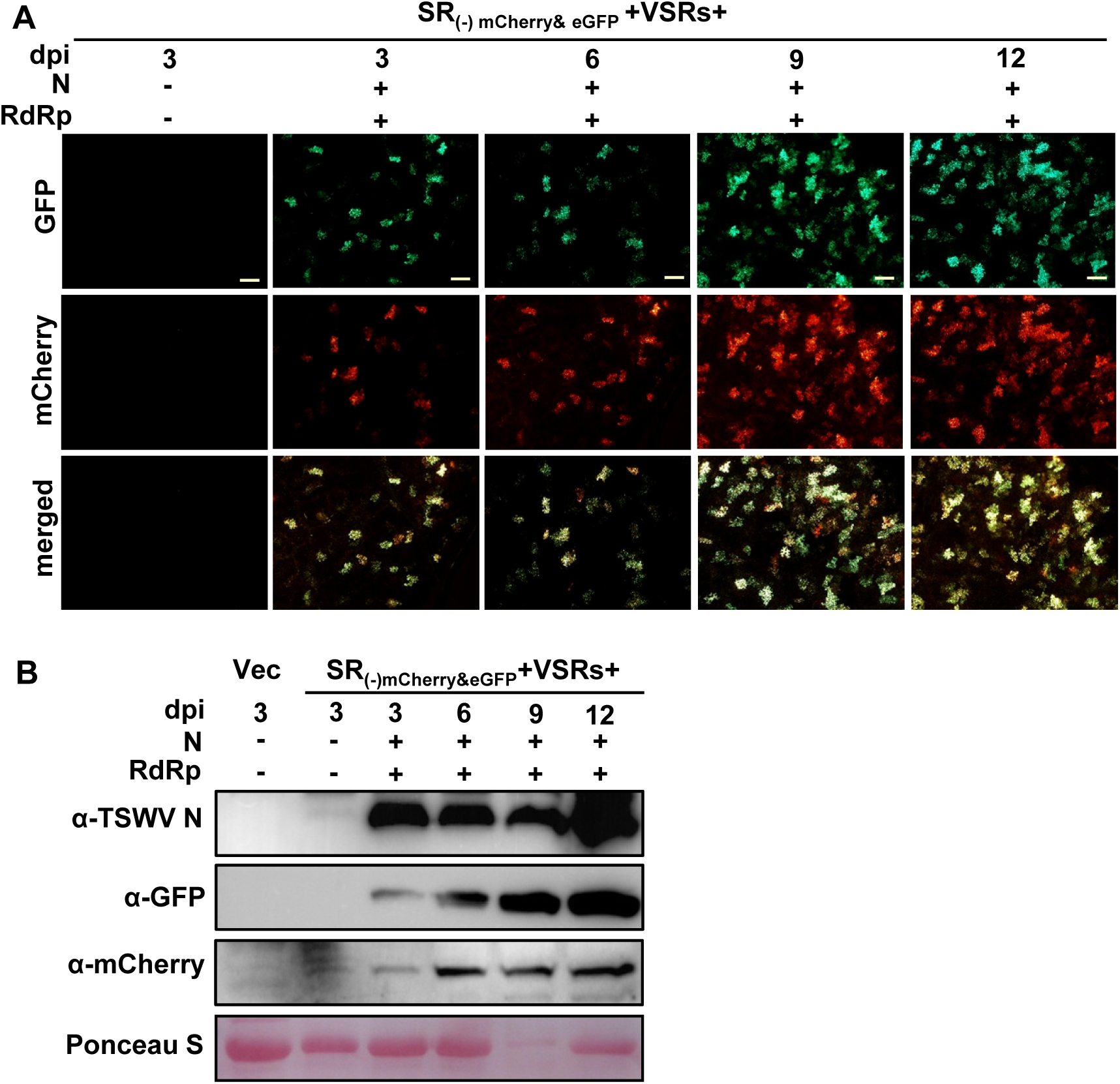
Time course analysis on gene expression from the SR_(-)mCherry&eGFP_ mini-replicon in *N. benthamiana* leaves. (*A*) Foci of eGFP and mCherry fluorescence expressed from SR_(-)mCherry&eGFP_ in *N. benthamiana* leaves co-expressing N, RdRp and the VSRs at 3, 6, 9 and 12 dpi, respectively. Fluorescence of eGFP and mCherry were photographed under a fluorescence microscope using GFP and RFP filters, respectively. Bars represent 400 μm. (*B*) Western immunoblot detection of the N, eGFP and mCherry proteins in leaves shown in panel A, using specific antibodies against N, GFP and mCherry, respectively. The empty vector (Vec) was used as a negative control. Ponceau S staining was used as protein loading control.

**Fig. S3.**
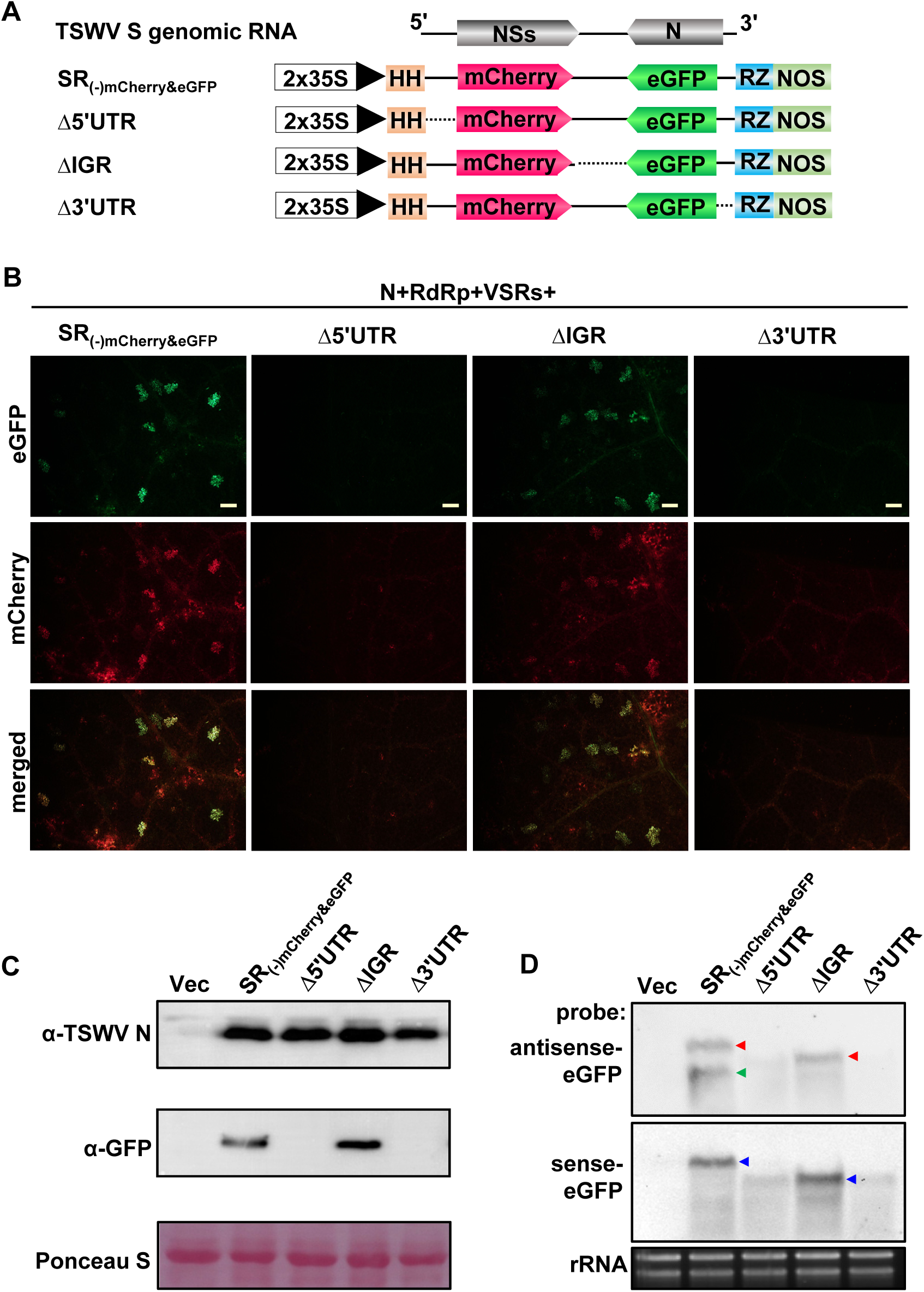
The role of 5’-UTR, 3’-UTR and IGR on viral RNA synthesis from the SR_(-)mCherry&eGFP_ mini-replicon. (*A*) Schematic representation of TSWV SR_(-)mCherry&eGFP_ and derivatives with deletions of the 5’UTR, IGR or 3’UTR. (*B*) eGFP and mCherry fluorescence expressed from TSWV SR_(-)mCherry&eGFP_ and mutant derivatives in *N. benthamiana*. The SR_(-)mCherry&eGFP_ or its mutants were coexpressed with N, RdRp and the four VSRs in *N. benthamiana* leaves. The agroinfiltrated leaves were examined and photographed at 5 dpi under a fluorescence microscope using GFP and RFP filters, respectively. Bars represent 200 μm. (*C*) Western immunoblot detection of the N and eGFP proteins expressed from the SR_(-)mCherry&eGFP_ mini-replicon and mutant derivatives, using specific antibodies against N and GFP, respectively. Ponceau S staining was used as protein loading control. (*D*) Northern blot analysis of viral RNA synthesis from SR_(-)mCherry&eGFP_ and mutant derivatives. The anti-genomic RNAs (red arrow), genomic RNAs (blue arrow) and eGFP mRNA transcripts (green arrow) were detected with DIG–labeled sense eGFP or anti-sense eGFP probes. Ethidium bromide staining was used as RNA loading control.

**Fig. S4.**
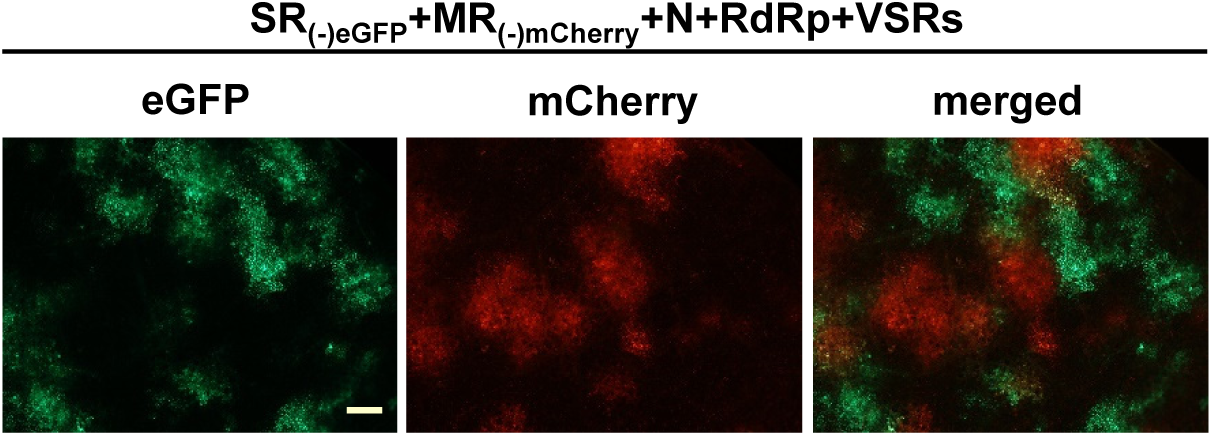
Cell-to-cell movement analysis of SR_(-)GFP_ and MR_(-)mCherry_ in *N. benthamiana* co-expressing RdRp, N and four VSRs in *N. benthamiana*. Agroinfiltrated leaves were examined and photographed at 5 dpi under a fluorescence microscope. Bars represent 400 μm.

**Fig. S5.**
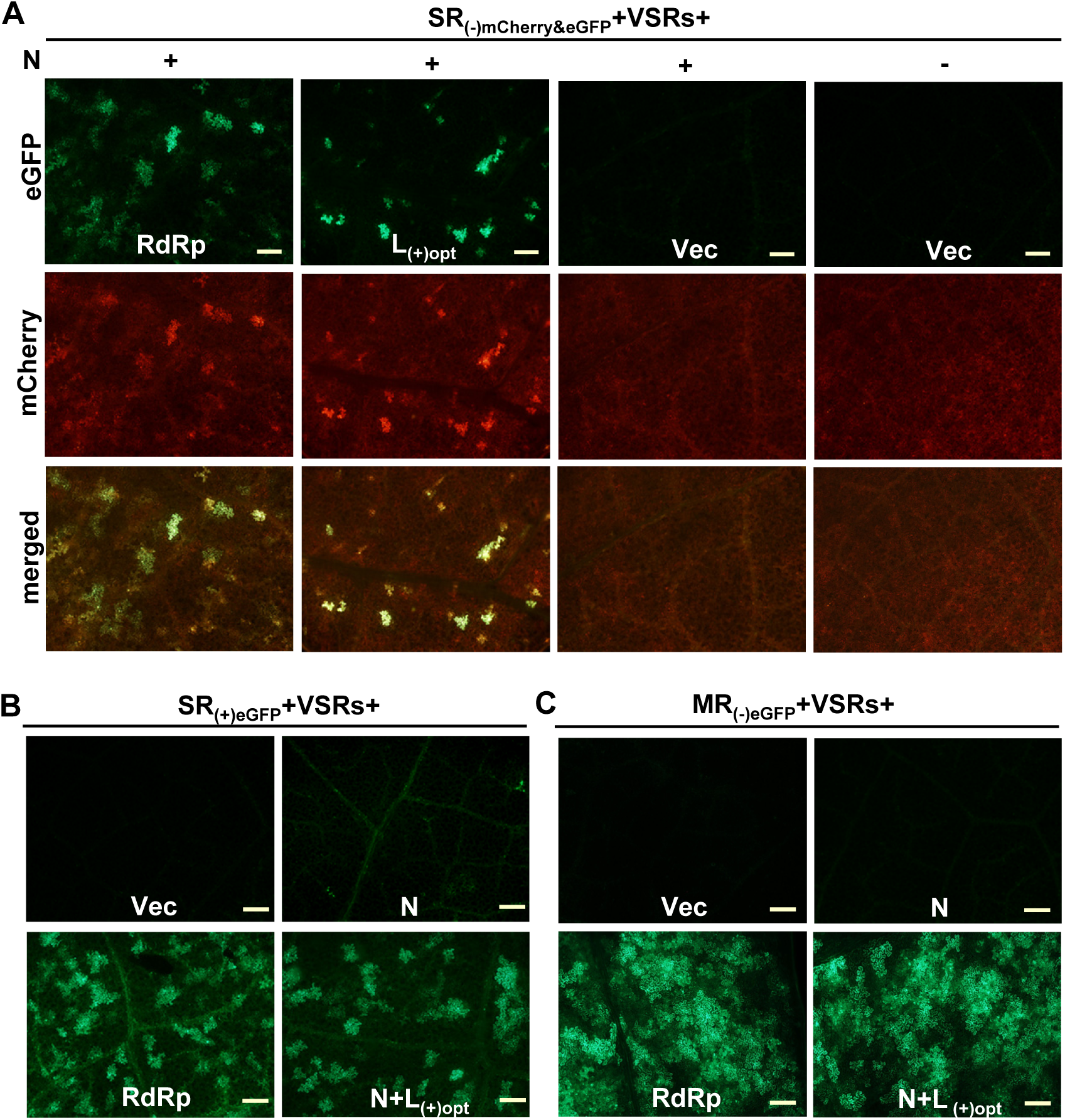
Functional analysis of RdRp expressed from TSWV L_(+)opt_, NSm from MR_(-)eGFP_ and N from SR_(+)eGFP_ using the mini-genome replication system in *N. benthamiana*. (*A*) Functional analysis of RdRp expressed from TSWV L_(+)opt_ using the S RNA mini-replicon system in *N. benthamiana*. The L_(+)opt_, RdRp, or pCB301 empty vector (Vec) was co-expressed with N, SR_(-)mCherry&eGFP_ and the four VSRs in *N. benthamiana* leaves. (*B*) Functional analysis of N expressed from SR_(+)eGFP_ in *N. benthamiana.* SR_(+)eGFP_ was co-expressed with the empty vector (Vec), N, RdRp or N+L_(+)opt_ in *N. benthamiana* leaves in the presence of four VSRs. (*C*) Functional analysis of NSm expressed from MR_(-)eGFP_ in *N. benthamiana.* MR_(-)eGFP_ was co-expressed with the empty vector (Vec), N, RdRp or N+L_(+)opt_ in *N. benthamiana* leaves in the presence of four VSRs. Foci showing mCherry and eGFP fluorescence in agroinfiltrated *N. benthamiana* leaves were examined at 3 dpi by a fluorescence microscope. Bars represent 400 μm.

**Fig. S6.**
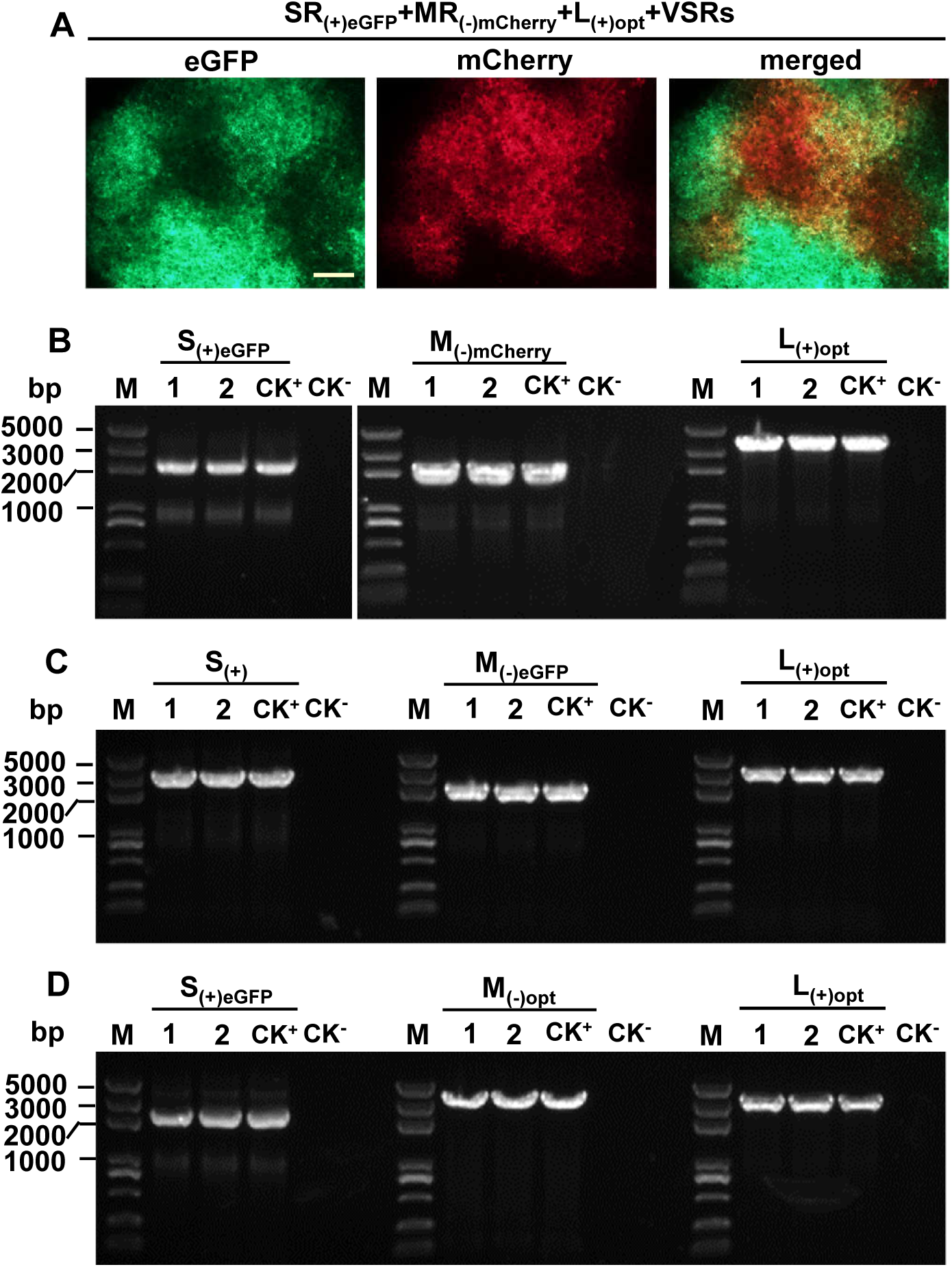
Analysis of *N. benthamiana* leaves agroinfiltrated with constructs of SR_(+)eGFP_, MR_(-)mCherry_ and L_(+)opt_, or SR_(+)eGFP_, MR_(-)opt_ and L_(+)opt_ or S_(+)_, MR_(-)eGFP_ and L_(+)opt_. (*A*) Local infection analysis of cell-to-cell movement of SR_(+)eGFP_ and MR_(-)mCherry_ co-expressing with L_(+)opt_ and four VSRs in *N. benthamiana* by agroinfiltration. The agro-infiltrated leaves were examined and photographed at 5 dpi under a fluorescence microscope. Bars represent 400 μm. (*B*) RT-PCR analysis on systemically infected leaves from *N. benthamiana* plants agroinfiltrated with SR_(+)eGFP_, MR_(-)mCherry_ and L_(+)opt_. (*C*) RT-PCR analysis on systemically infected leaves from *N. benthamiana* plants agroinfiltrated with S_(+)_, MR_(-)eGFP_ and L_(+)opt_ (*D*) RT-PCR analysis on systemically infected leaves from *N. benthamiana* plants agroinfiltrated with SR_(+)eGFP_, M_(-)opt_ and L_(+)opt_. All agroinfiltrations were performed in the additional presence four VSRs constructs. RT PCR was performed on total RNA purified from systemic leaves for detection of S, M or L segments using segment-specific primers. Amplicons were resolved by electrophoresis in a 1 % agarose gel. Lanes 1-2 represent two biological replicates of systemic infected leaf samples; As positive controls (CK^+^) for proper fragment size, PCR was performed on plasmids carrying S, M, L_(+)opt_ or derivatives. As negative control (CK^-^), RT-PCR was performed in the absence of nucleic acids. DNA size markers are shown on the left hand side.

**Fig. S7.**
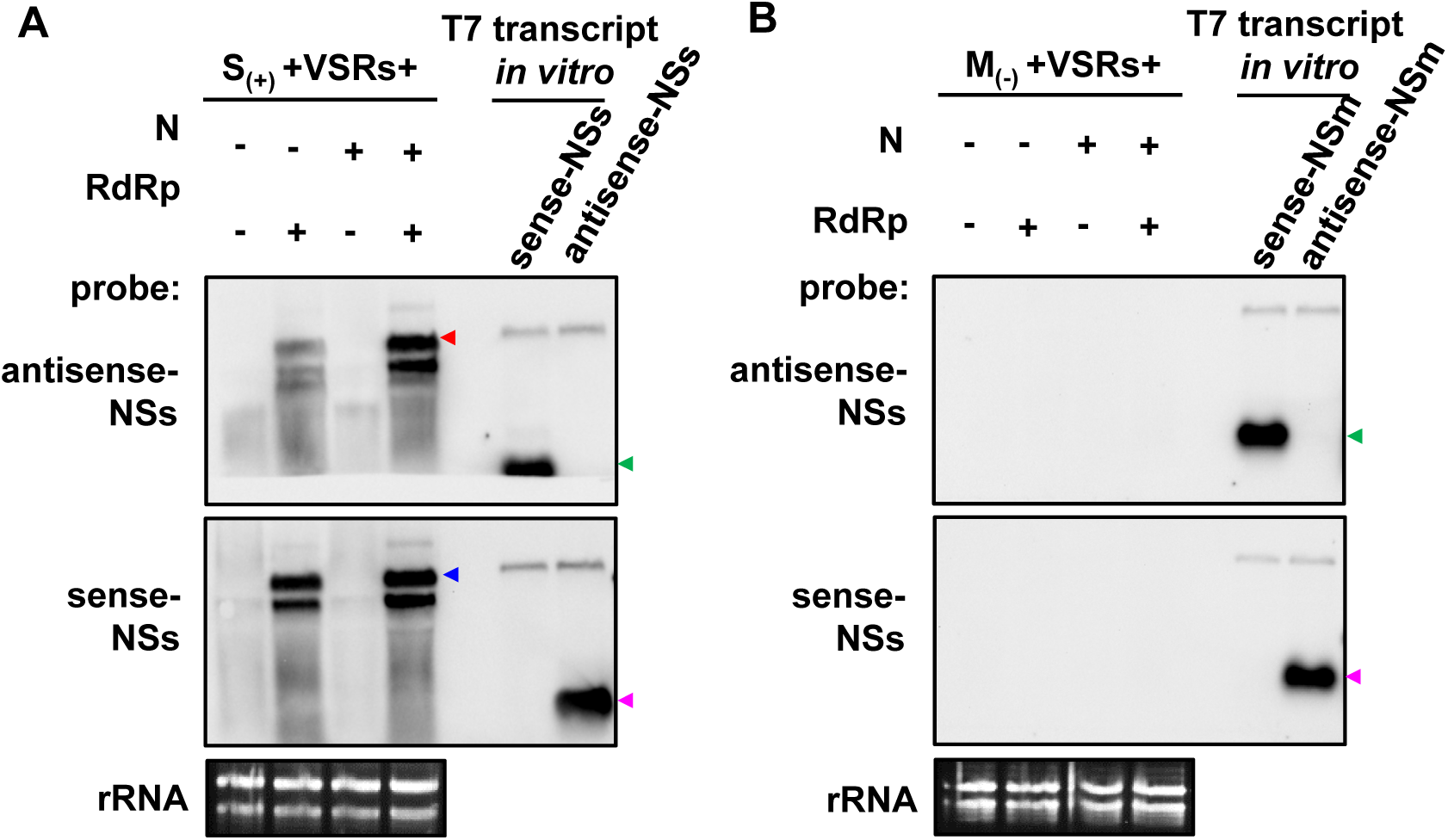
Northern blot detection of viral RNA synthesis produced from full length S_(+)_ and wild type M_(-)_ replicons. (*A* and *B*) The full length S_(+)_ or wild type M_(-)_ was co-expressed with the empty vector (Vec), N, RdRp_opt_ or N+RdRp_opt_ in *N. benthamiana* leaves in the presence of four VSRs. The genomic RNAs (blue arrow), anti-genomic RNAs (red arrow) of S (*A*) and M (*B*) were detected with DIG–labeled sense or anti-sense NSs and NSm probes, respectively. Ethidium bromide staining was used as RNA loading control.

**Fig. S8.**
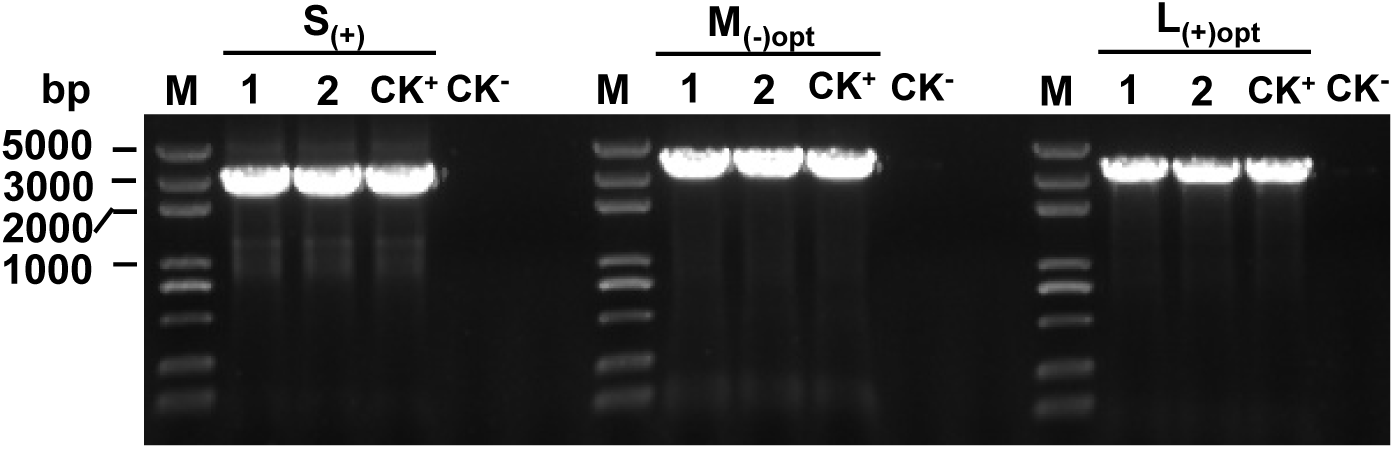
RT-PCR detection of S_(+)_, M_(-)opt_ and L_(+)opt_ genomic RNA in systemic leaves of *N. benthamiana* infected by rTSWV. The S_(+)_, M_(-)opt_ and L_(+)opt_ and the four VSRs were co-expressed in *N. benthamiana* leaves by agroinfiltration. Total RNA was purified from systemic leaves of agroinfiltrated plants and the presence of S_(+)_, M_(-)opt_ and L_(+)opt_ were detected by RT-PCR using segment-specific primers. RT-PCR products were resolved by electrophoresis in a 1% agarose gel. Lanes 1-2, two biological replicates of systemic infected leaf samples; RT-PCR on plasmids carrying S, M and L as DNA template were used as positive controls (CK^+^). RT-PCR without adding the DNA template was used as negative controls (CK^-^). DNA size markers are shown on the left hand side.

**Fig. S9.**
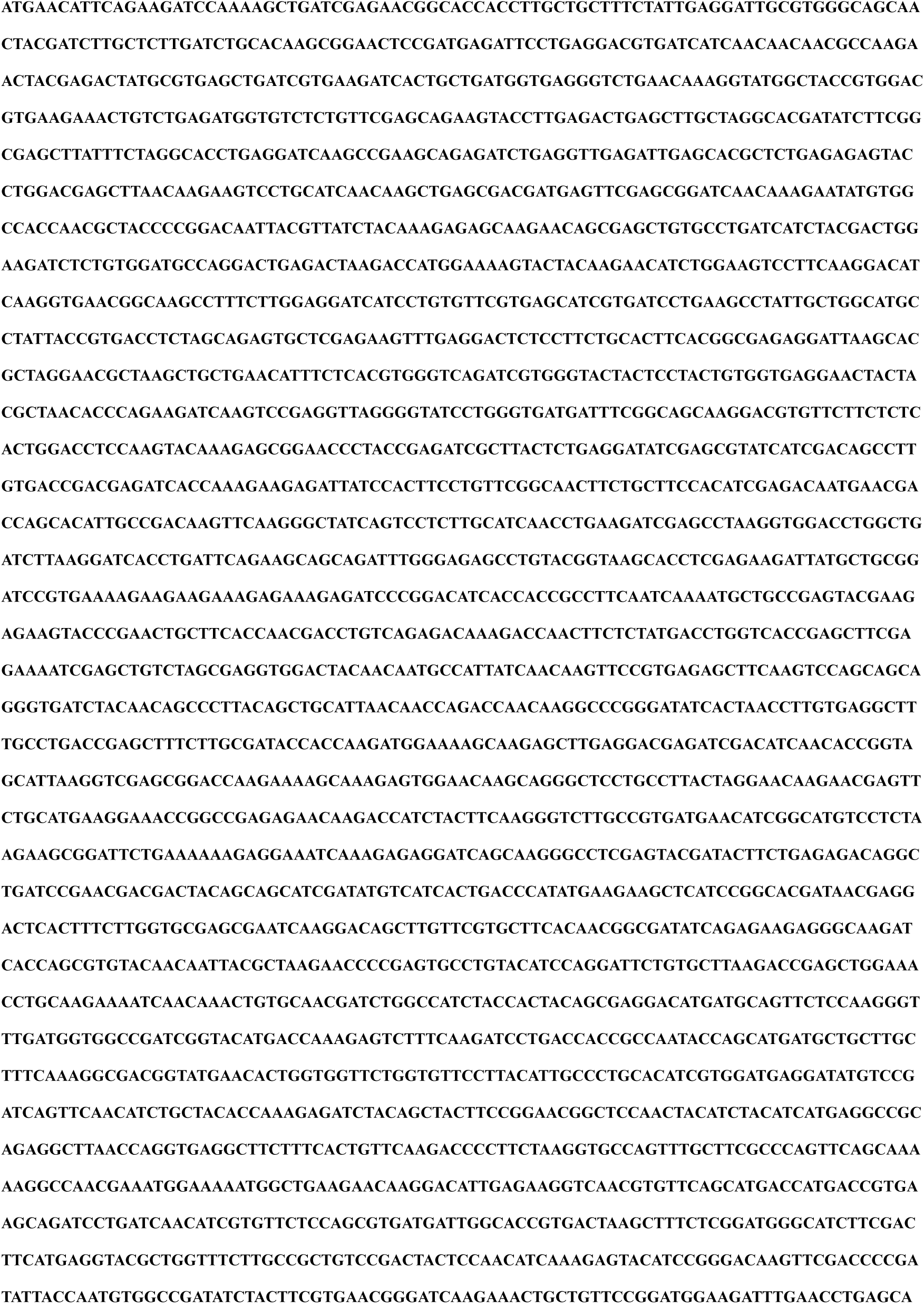
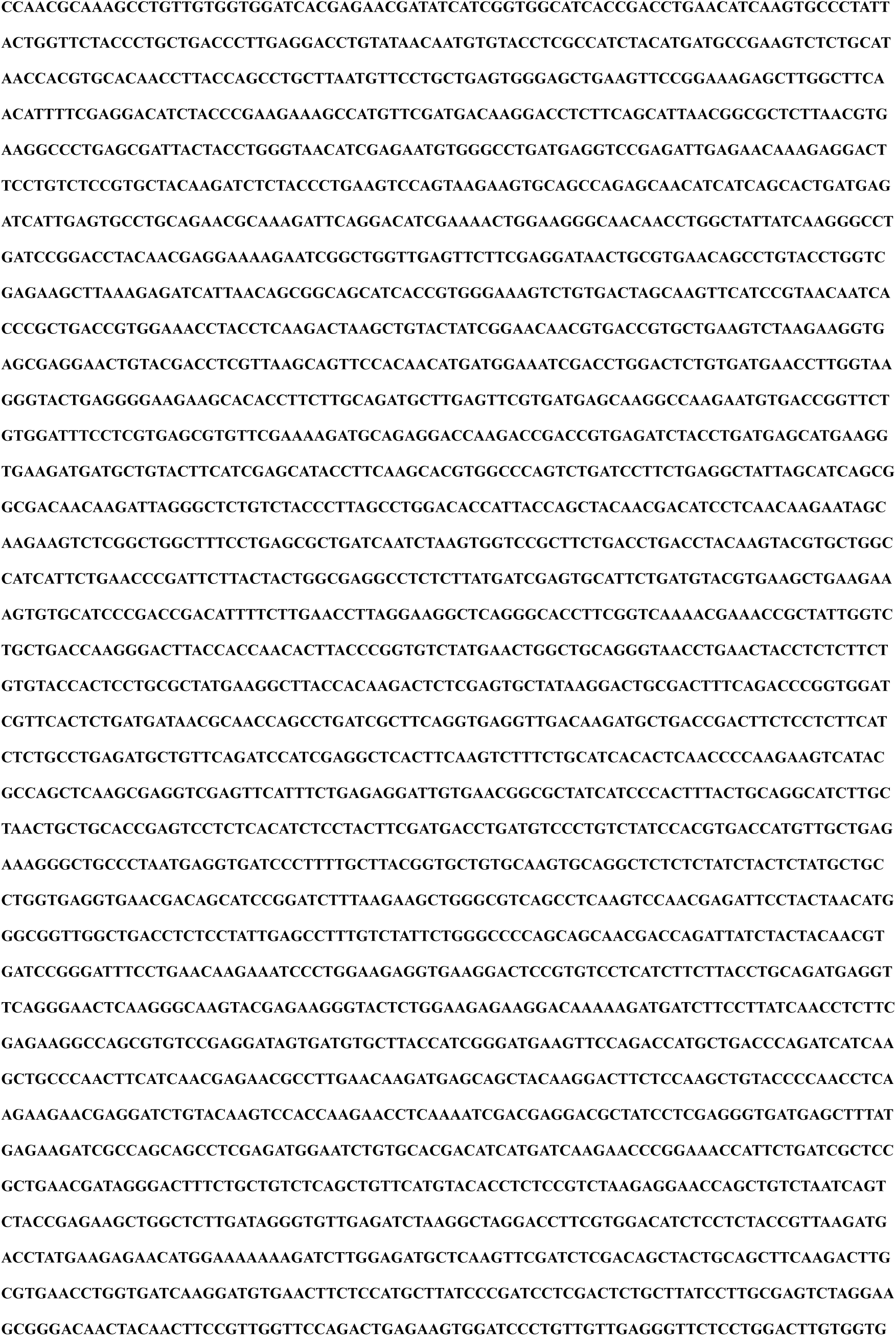
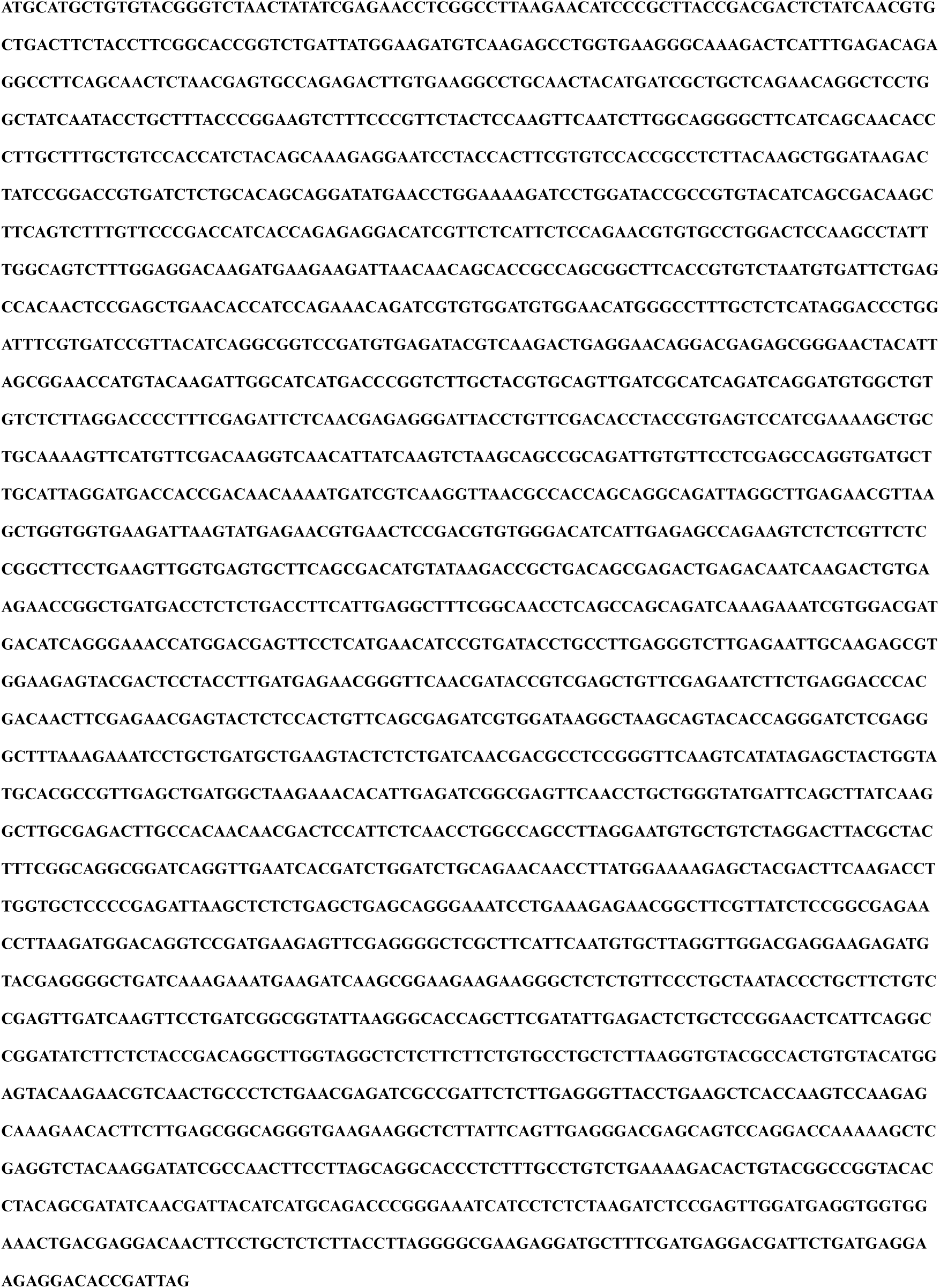
Optimized RdRp gene sequence used in the study.

**Fig. S10.**
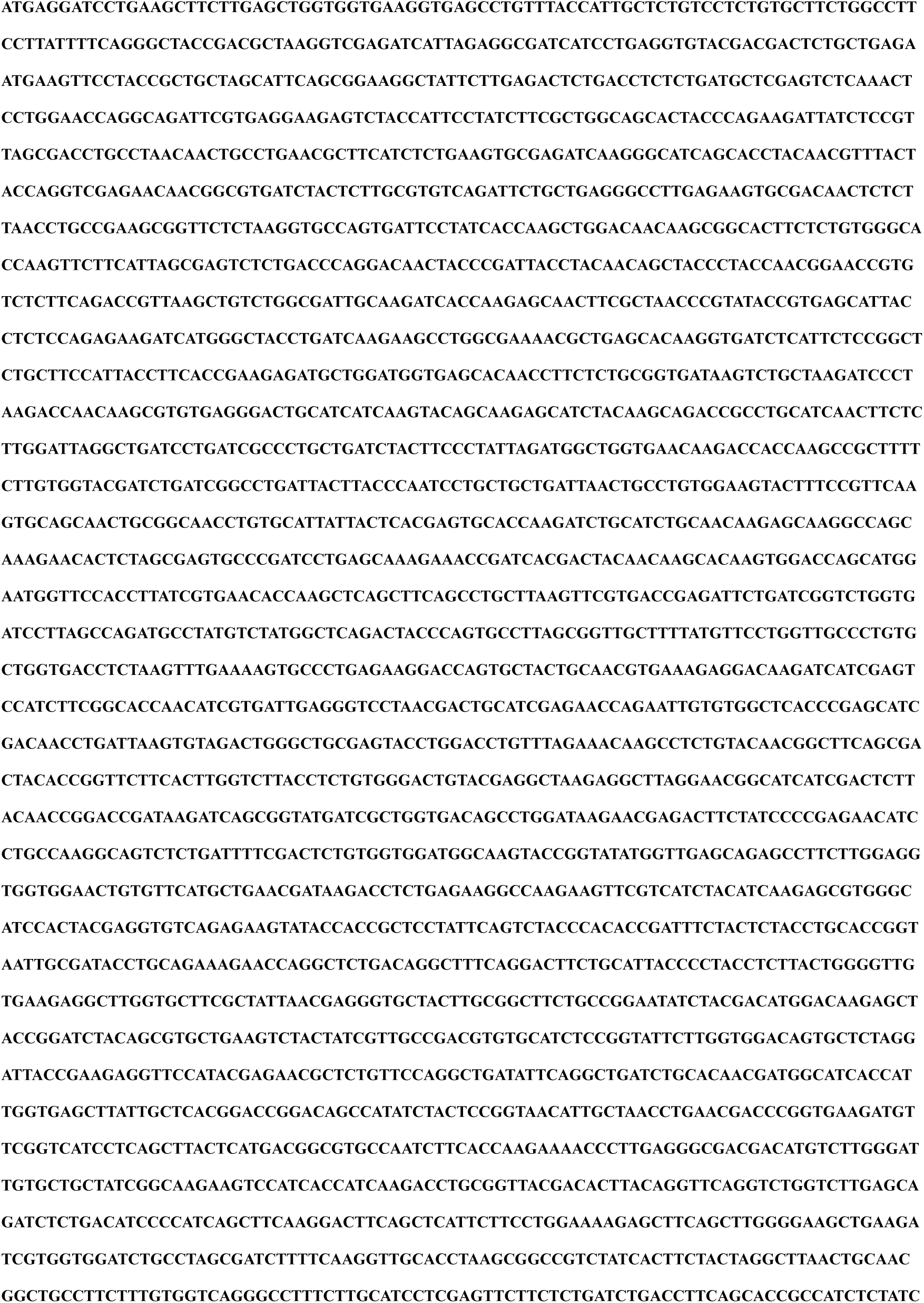
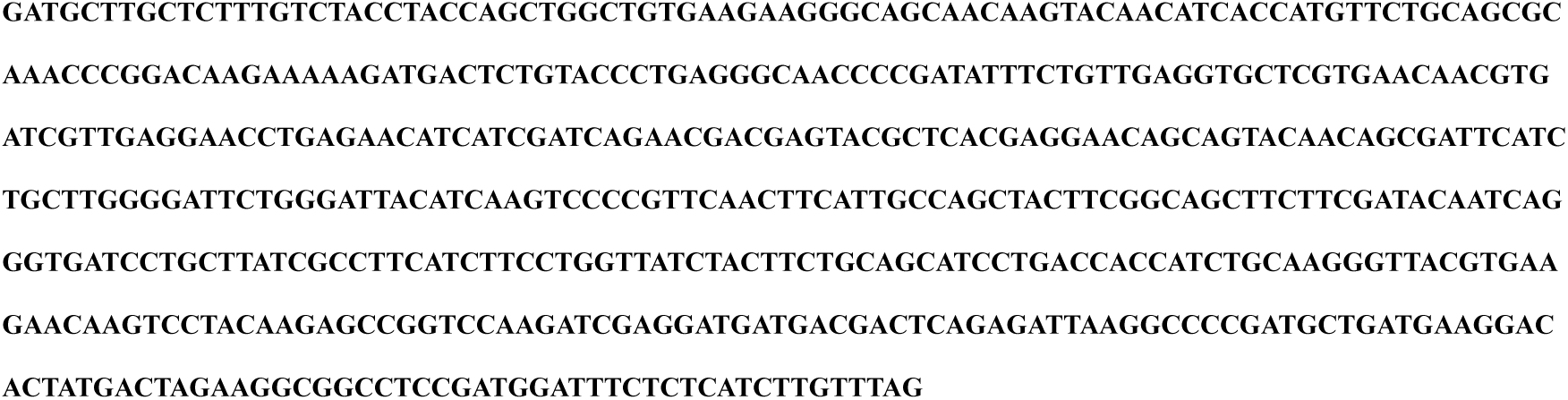
Optimized GP gene sequence used in the study.

## Supplementary Materials and Methods

### Plasmid construction

#### Construction of RdRp, RdRp_opt_, N, NSs and VSRs

The cDNA of the RdRp, N, and NSs genes was amplified from the total RNA of TSWV-lettuce isolate infected tissues and inserted into a binary vector pCambia2300 or pCXSN to generate p2300-RdRp_wt_, p2300-N, and pCXSN-NSs downstream a double 35S promoter (2×35S). The P19-HcPro-γb carrying three VSRs simultaneously in the pCB301 vector was kindly provided by Dr. Xianbing Wang in College of Biological Sciences of China Agricultural University. The codon usage and intron-splicing sites optimized sequence of RdRp (*SI Appendix,* Fig. S9) was *de novo* synthesized by GenScript Biotech Corp (Nanjing, China) and was inserted into a binary vector pCambia2300 to generate p2300S-RdRp_opt_ downstream a 2×35S promoter.

#### Construction of full length TSWV genomic S_(-)_, M_(-)_, L_(-)_, and anti-genomic S_(+)_, M_(+)_ and L_(+)_ cDNA clones

To generate constructs to express full length TSWV genomic RNA and antigenomic RNA S, M and L segments, total RNA extracted from TSWV-lettuce infected leaves of *N. benthamiana* plants was reverse transcribed into cDNA, followed by PCR amplification with specific primers (*SI Appendix,* Table S3) using Phanta Super-Fidelity DNA Polymerase (Vazyme Biotech, Nanjing, China). The PCR products were fused with a self-cleaving hammerhead (HH) ribozyme (1) and inserted into a binary expression vector pCB301-2×35S-RZ-NOS linearized by two restriction endonucleases *Stu* I and *Sma* I (2). The pCB301-2×35S-HH-S_(-)_-RZ-NOS [S_(-)_], pCB301-2×35S-HH-M_(-)_-RZ-NOS [M_(-)_], pCB301-2×35S-HH-L_(-)_-RZ-NOS [L_(-)_], pCB301-pCB301-2×35S-HH-S_(+)_-RZ-NOS [S_(+)_], 2×35S-HH-M_(+)_-RZ-NOS [M_(+)_] and pCB301-2×35S-HH-L_(+)_-RZ-NOS [L_(+)_] cDNA clones were generated. The full length TSWV genomic S_(-)_, M_(-)_ and L_(-)_, and anti-genomic S_(+)_, M_(+)_ and L_(+)_ were expressed downstream a double 35S promoter (2×35S) and franked with a self-cleaving hammerhead (HH) ribozyme at 5’-terminus and a Hepatitis delta virus (HDV) ribozyme at 3’-terminus.

#### Construction of TSWV SR_(-)eGFP_, SR_(-)mCherry&eGFP_ and SR_(+)eGFP_ minireplicons

To generate SR_(-)eGFP_-genomic RNA minireplicon, the eGFP ORF was amplified and used to replace the N gene in the pCB301-2×35S-HH-S_(-)_-RZ-NOS by the *in vitro* recombination using the In-Fusion Cloning mixture (Clontech, Japan). The construct pCB301-2×35S-HH-SR_(-)eGFP_-RZ-NOS [SR_(-)eGFP_] was generated.

To generate SR_(-)mCherry&eGFP_ in which the NSs and N genes in S gRNA were replaced with mCherry and eGFP, respectively, the mCherry ORF was amplified and used to exchange the NSs gene in the pCB301-2×35S-HH-SR_(-)eGFP_-RZ-NOS by recombination using In-Fusion Cloning mixture (Clontech). The construct pCB301-2×35S-HH-SR_(-)mCherry&eGFP_-RZ-NOS [35S:SR_(-))mCherry&eGFP_] was generated. The T7:SR_(-)mCherry&eGFP_ minireplicon (pCB301-T7-HH-SR_(-)mCherry&eGFP_-RZ-NOS) controlled by T7 promoter was constructed by the same strategy as 35S:SR_(-)mCherry&eGFP_.

To generate antigenomic S_(+)eGFP_-minireplicon, the eGFP ORF was amplified and used to replace the NSs gene in the pCB301-2×35S-HH-S_(-)_-RZ-NOS by recombination using In-Fusion Cloning mixture (Clontech). The construct pCB301-2×35S-HH-SR_(+)eGFP_-RZ-NOS [SR_(+)eGFP_] was generated. The primers used above are listed in *SI Appendix,* Table S3.

#### Construction of SR_(-)mCherry&eGFPΔ5’UTR_, SR_(-)mCherry&eGFPΔIGR_ and SR_(-)mCherry&eGFPΔ3’UTR_ mutants

To generate SR_(-)mCherry&eGFP_**_Δ_**_5’UTR_, the DNA copy of SR_(-)mCherry&eGFP_ without 5’-UTR (88 nt) was amplified from pCB301-2×35S-HH-SR_(-)mCherry&eGFP_-RZ-NOS and used to recombine with backbone vector of pCB301-2×35S-HH-SR_(-)mCherry&eGFP_-RZ-NOS by recombination using In-Fusion Cloning mixture (Clontech). The construct pCB301-2×35S-HH-SR_(-)mCherry&eGFP**Δ**5’UTR_-RZ-NOS [SR_(-)mCherry&eGFP**Δ**5’UTR_] was generated.

To generate SR_(-)mCherry&eGFP**Δ**IGR_, the DNA copy of SR_(-)mCherry&eGFP_ without IGR (550 nt) was amplified from pCB301-2×35S-HH-SR_(-)mCherry&eGFP_-RZ-NOS and used to recombine with the vector backbone from pCB301-2×35S-HH-SR_(-)mCherry&eGFP_-RZ-NOS by recombination using In-Fusion Cloning mixture (Clontech). The construct pCB301-2×35S-HH-SR_(-)mCherry&eGFP**Δ**IGR_-RZ-NOS [SR_(-)mCherry&eGFP**Δ**IGR_] was generated.

To generate SR_(-)mCherry&eGFP_**_Δ_**_3’UTR_, the DNA copy of SR_(-)mCherry&eGFP_ without 3’-UTR (151 nt) was amplified from pCB301-2×35S-HH-SR_(-)mCherry&eGFP_-RZ-NOS and used to recombine with the vector backbone from pCB301-2×35S-HH-SR_(-)mCherry&eGFP_-RZ-NOS using the primer pair FMF48/P3382 by recombination using In-Fusion Cloning mixture (Clontech). The construct pCB301-2×35S-HH-SR_(-)mCherry&eGFP_**_Δ_**_3’UTR_-RZ-NOS [SR_(-)mCherry&eGFP**Δ**3’UTR_] was generated. The primers used above are listed in *SI Appendix,* Table S3.

#### Construction of TSWV MR_(-)eGFP_, MR_(-)mCherry_ and MR_(-)eGFPNSmMut_ minireplicons

To generate MR_(-)eGFP_ and MR_(-)mCherry_ minireplicons, the eGFP and mCherry ORFs were amplified and used to replace the GP gene in pCB3012×35S-HH-M_(-)_-RZ-NOS, respectively, by recombination using In-Fusion Cloning mixture (Clontech). The constructs pCB301-2×35S-HH-MR_(-)eGFP_-RZ-NOS [MR_(-)eGFP_] and pCB301-2×35S-HH-MR_(-)mCherry_-RZ-NOS [MR_(-)mCherry_] was generated.

To generate MR_(-)eGFPNSmMut_ in which a stop codon was introduced immediately after the start codon of NSm, the NSm_Mut_ was amplified and used to replace the wild type NSm sequence in pCB301-2×35S-HH-MR_(-)eGFP_-RZ-NOS by recombination using In-Fusion Cloning mixture (Clontech). The construct pCB301-2×35S-HH-MR_(-)eGFPNSmMut_-RZ-NOS [MR_(-)eGFPNSmMut_] was generated. All primers used above are listed in *SI Appendix,* Table S3.

#### Construction of full length L_(+)opt_ and M_(-)opt_ cDNA clones

To generate full length L_(+)opt_ cDNA clone, the sequence codon and intron-splicing sites optimized RdRp was amplified and used to replace the wild type RdRp sequence in pCB301-2×35S-HH-L_(+)_-RZ-NOS by recombination using the In-Fusion Cloning mixture (Clontech). The pCB301-2×35S-HH-L_(-)opt_-RZ-NOS [L_(-)opt_] was generated.

To generate full length M_(-)opt_ cDNA clone, the codon and intron-splicing sites optimized GP gene was de novo synthesized by GenScript Biotech Corp (Nanjing, China) (*SI Appendix,* Fig. S10) and used to replace the wild type GP sequence in pCB301-2×35S-HH-M_(-)_-RZ-NOS by the *in vitro* recombination using In-Fusion Cloning mixture (Clontech). The pCB301-2×35S-HH-M_(-)opt_-RZ-NOS [M_(-)opt_] was generated. The primers used above are listed in *SI Appendix,* Table S3.

### Plant material and virus source

Six to eight weeks of *Nicotiana benthamiana* was used in all agroinfiltration assay. *N. benthamiana* plants were grown in a growth chamber setting at 25 °C, a 16 h light and 8 h dark photoperiod (3). The TSWV isolate from asparagus lettuce (TSWV-LE) was used in this study (GenBank accession number: KU976396 for S, JN664253 for M and KU976394 for L) (4). The TSWV-LE isolate was maintained on *N. benthamiana*. For long-term storage, the infected new leaves of *N. benthamiana* were kept in an 80 °C refrigerator.

### Agrobacterium infiltration

Recombinant plasmids were electroporated into *Agrobacterium tumefaciens* strain GV3101 and agroinfiltrations were performed essentially as described (5, 6). *A. tumefaciens* cells were resuspended by agroinfiltration buffer [10 mM MgCl2, 10 mM MES (pH 5.6) and 100 μM acetosyringone] adjusted to an optical density OD_600_ of 1.0 and incubated for 2 to 3 h in dark at room temperature. Equal volumes of *Agrobacterium* cultures (final concentration OD_600_=0.2) harboring the p2300-N, p2300-RdRp, pCB301-derived reporter or full-length infectious clone vector(s), were mixed with one volume of bacterial mixture (final concentration OD_600_=0.05) containing the NSs and P19-HcPro-γb. The *Agrobacterium* cultures were infiltrated into fully expanded leaves of 6-7 leaf stage *N. benthaminan* plants using 1 mL needleless syringes.

### Immunoblot analysis

Total protein was extracted from 1 g *Agrobacterium*-infiltrated leaf patches, healthy or TSWV-infected *N. benthamiana* systemic leaves in a 1 mL extraction buffer [10 % (v/v) glycerol, 25 mM Tris-HCl, pH 7.5, 1 mM EDTA, 150 mM NaCl, 10 mM dithiothreitol, 2 % (w/v) polyvinylpolypyrrolidone, 0.5 % (v/v) Triton X-100 and 1× protease inhibitors cocktail] (7). Protein samples were separated by SDS-PAGE gels, transferred to PVDF membranes (GE Healthcare, UK), blocked with 5 % skim milk solution and incubated with a polyclonal antiserum specific to the TSWV N, NSm, NSs, Gn, Gc, GFP, mCherry or T7 RNA pol at room temperature for 1 h or overnight at 4 °C and washed three times. After incubation in a secondary antibody containing HRP-conjugated goat anti-rabbit (1:10000) for 1 h, the blots were detected using the ECL Substrate Kit (Thermo Scientific, Hudson, NH, USA). To evaluate protein loading, the blots were stained with Ponceau S.

### Northern blot analysis

For Northern blot analysis of TSWV gRNAs, agRNAs or viral mRNA transcripts, total RNAs were extracted from *Agrobacterium*-infiltrated leaf patches, healthy or TSWV-infected systemic leaves using an RNAprep Pure Plant Kit (Tiangen Biotech, Beijing, China), respectively. DIG-labeled specific probes for sense or antisense GFP, NSs, NSm, L-5’UTR was synthesized by DIG High Prime RNA labeling kit (Roche, Basel, Switzerland). The total RNAs were separated on 1 % formaldehyde agarose gels and transferred to Hybond-N+ membranes (GE Healthcare, UK) (8). The membrane blots were hybridized with a DIG-labeled specific probe and detected using a DIG-High Prime Detection Starter Kit II (Roche), following the manufacturer’s protocol.

### RT-PCR and sequencing analysis

To detect the virus in systemic leaves of *N. benthamina* infected with SR_(+)eGFP_+MR_(-)mCherry_+L_(+)opt_, S_(+)_+MR_(-)eGFP_+L_(+)opt_, SR_(+)eGFP_+M_(-)opt_+L_(+)opt_ or rTSWV recovered from the full-length cDNA clones, total RNAs were extracted from systemic symptoms plant leaves. First-strand cDNAs were synthesized using M-MLV Reverse Transcriptase (Promega, USA). RT-PCRs were performed to detect the SR_(+)eGFP_, MR_(-)mCherry_, MR_(-)eGFP_, S_(+)_, M_(-)opt_ and L_(+)opt_ minigenome and genomic RNA using their specific-primers. The PCR products were inserted into a pMD19-T vector (Takara, Dalian, China) and sequenced by Sanger dideoxy-mediated chain-termination DNA sequencing method at Sangon Biotech (Shanghai, China). The primers used in this study are listed in *SI Appendix,* Table S3.

### Fluorescence microscopy

The agro-infiltrated *N. benthamiana* leaves were examined for fluorescence expression using an OLYMPUS IX71-F22FL/DIC Inverted Fluorescence Microscope (OLYMPUS, Tokyo, Japan) with a green or red barrier filter. The leaf sample was fixed in water on a microslider under a coverslip to detect the eGFP and mCherry fluorescence, respectively. Fluorescence images were processed using ImagePro (OLYMPUS, Tokyo, Japan) and Adobe (San Jose, CA, USA) Photoshop programs.

### Electron microscopy

Small tissues (1 mm × 4 mm) were excised from leaves of the *N. benthamiana* with infected rTSWV rescued from the full-length infectious clones. The sample tissues were fixed in 2.5 % glutaraldehyde and 1 % osmium tetroxide dissolving into 100 mM phosphate buffer (pH 7.0) as described by Li *et al* (5, 9) and then embedded in Epon 812 resin as instructed by the manufacture (SPI-EM, Division of Structure Probe, Inc., West Chester, USA). Ultrathin sections (70 nm) were mounted on formvar-coate grids and then stained with uranyl acetate for 10 min followed by lead citrate for 10 min. The stained sections were examined under a transmission electron microscope (TEM; H-7650, Hitachi, Japan).

### Imaging GFP in infected plant by hand-held UV lamp

GFP fluorescence in leaves was monitored with a hand-held 100 W, long-wave UV lamp (UV Products, Upland, CA, USA) and the leaves were photographed using a Canon EOS 70D digital camera (Canon, Japan) with a 58 mm UV filter.

---

**Table S1.**
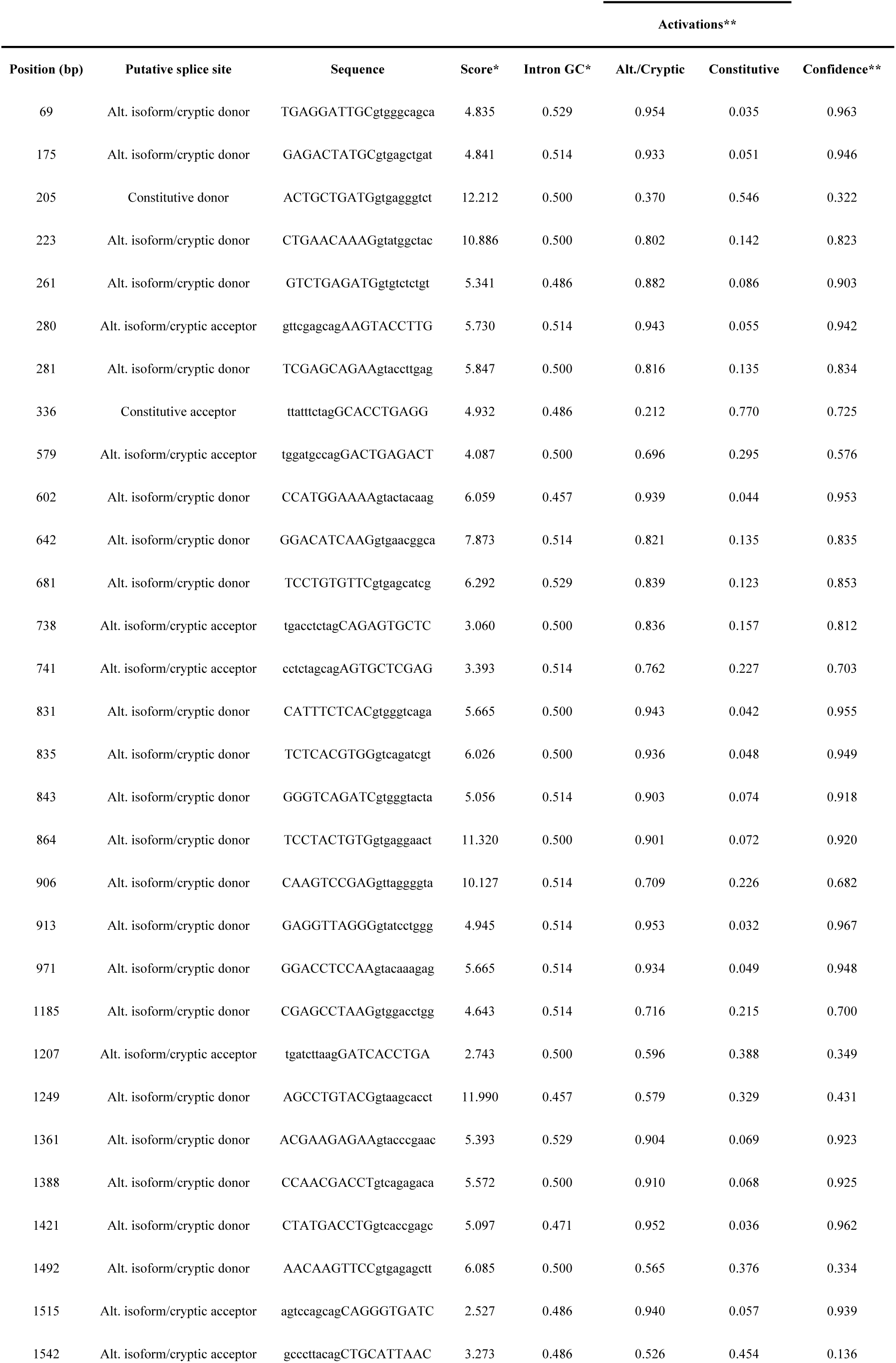
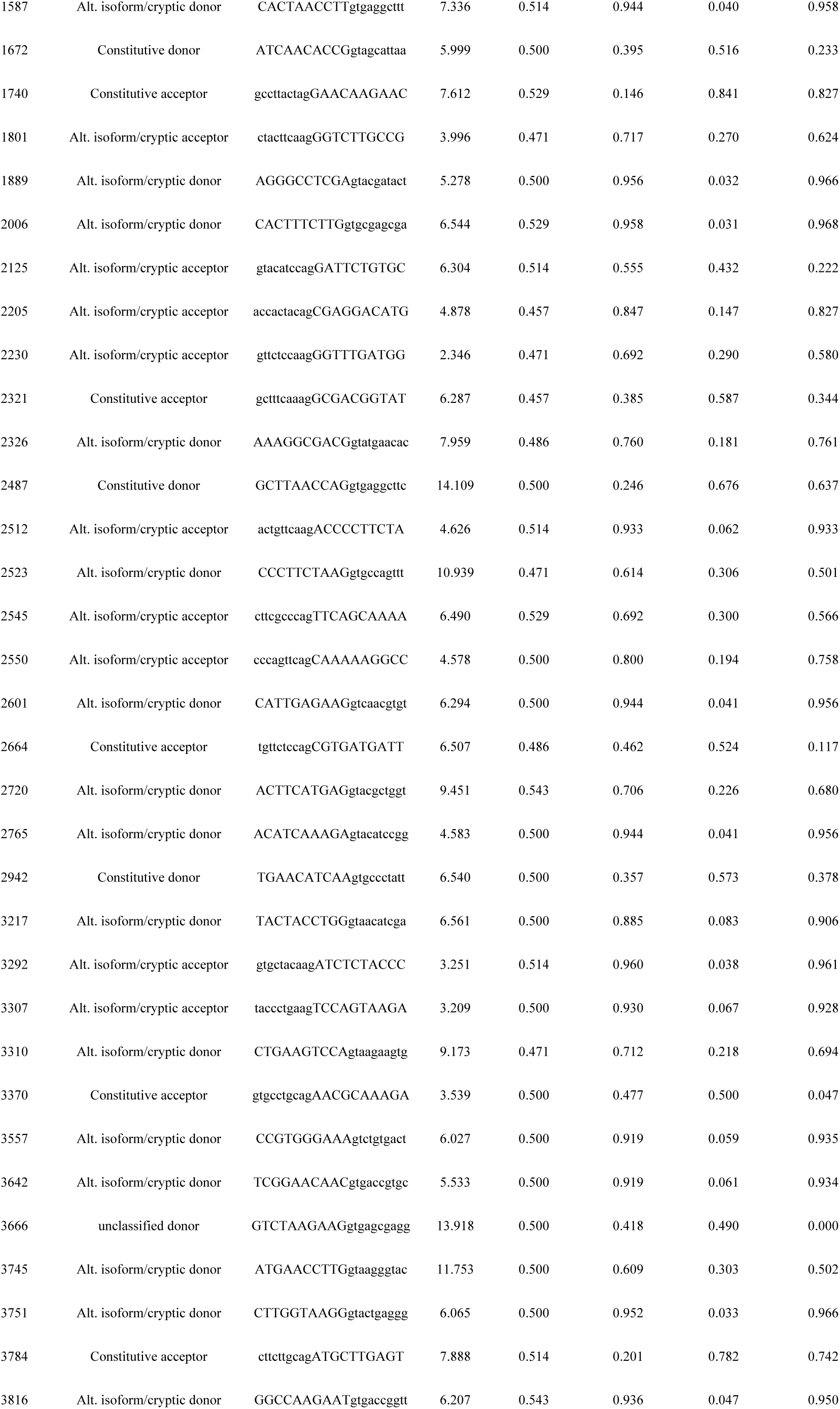
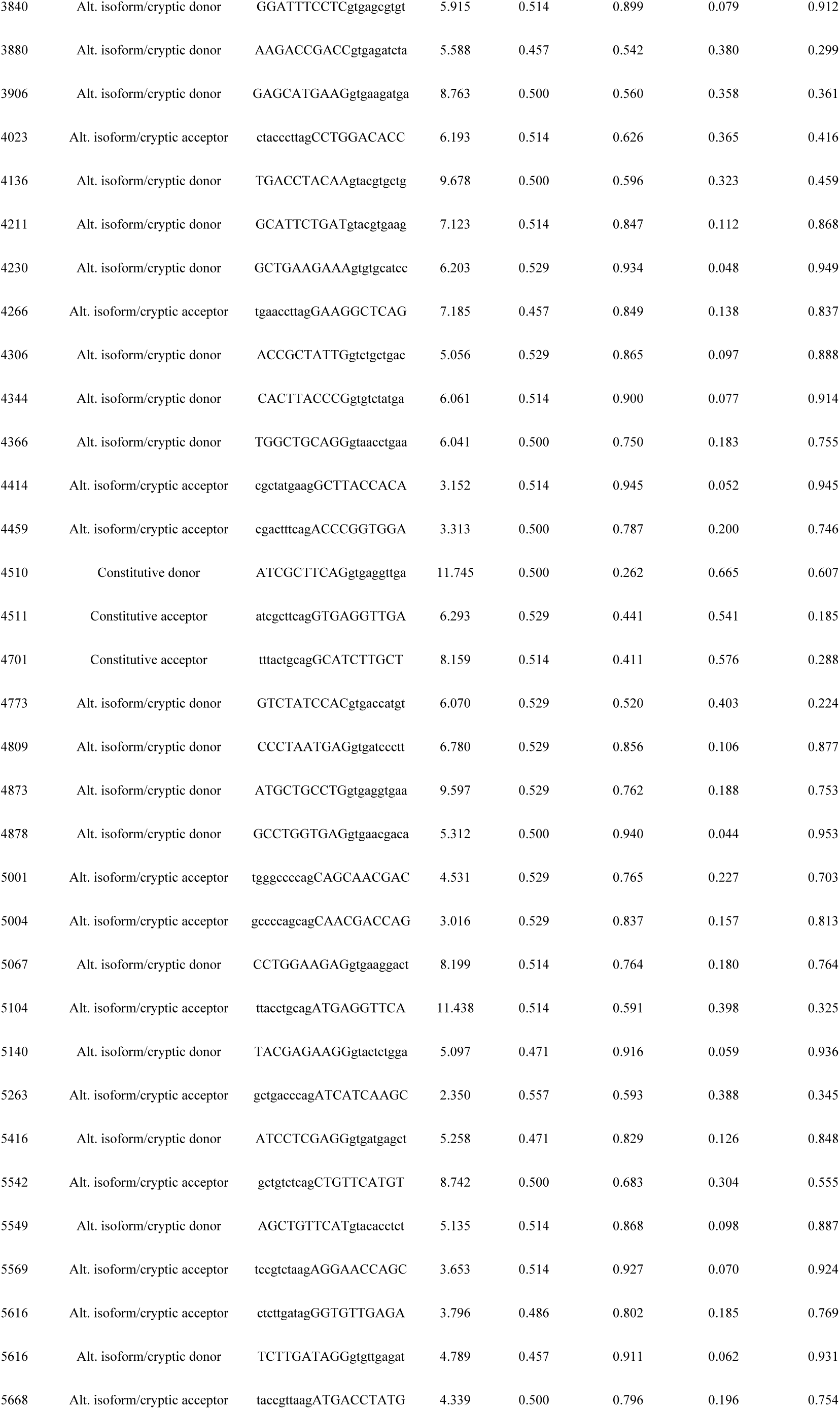
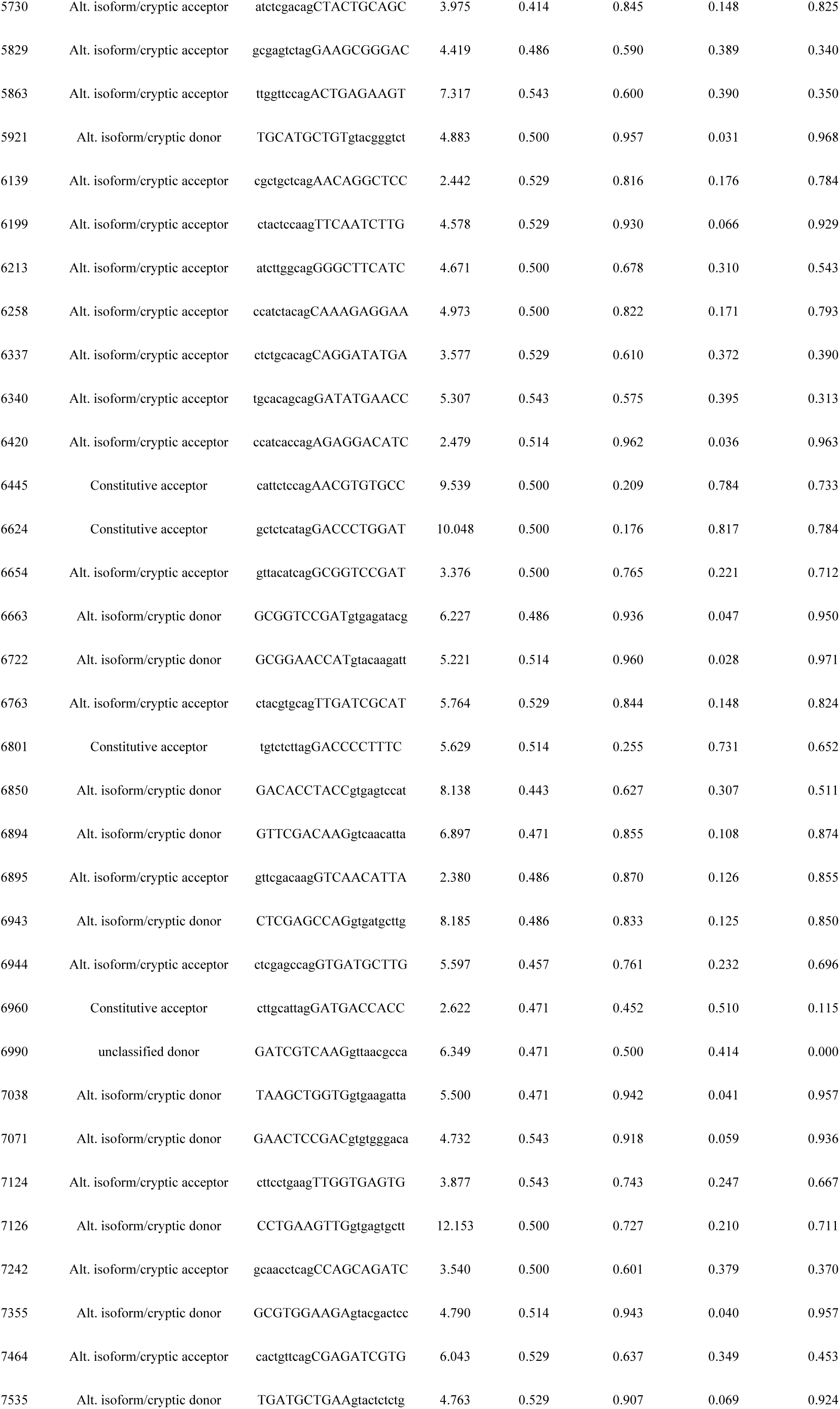
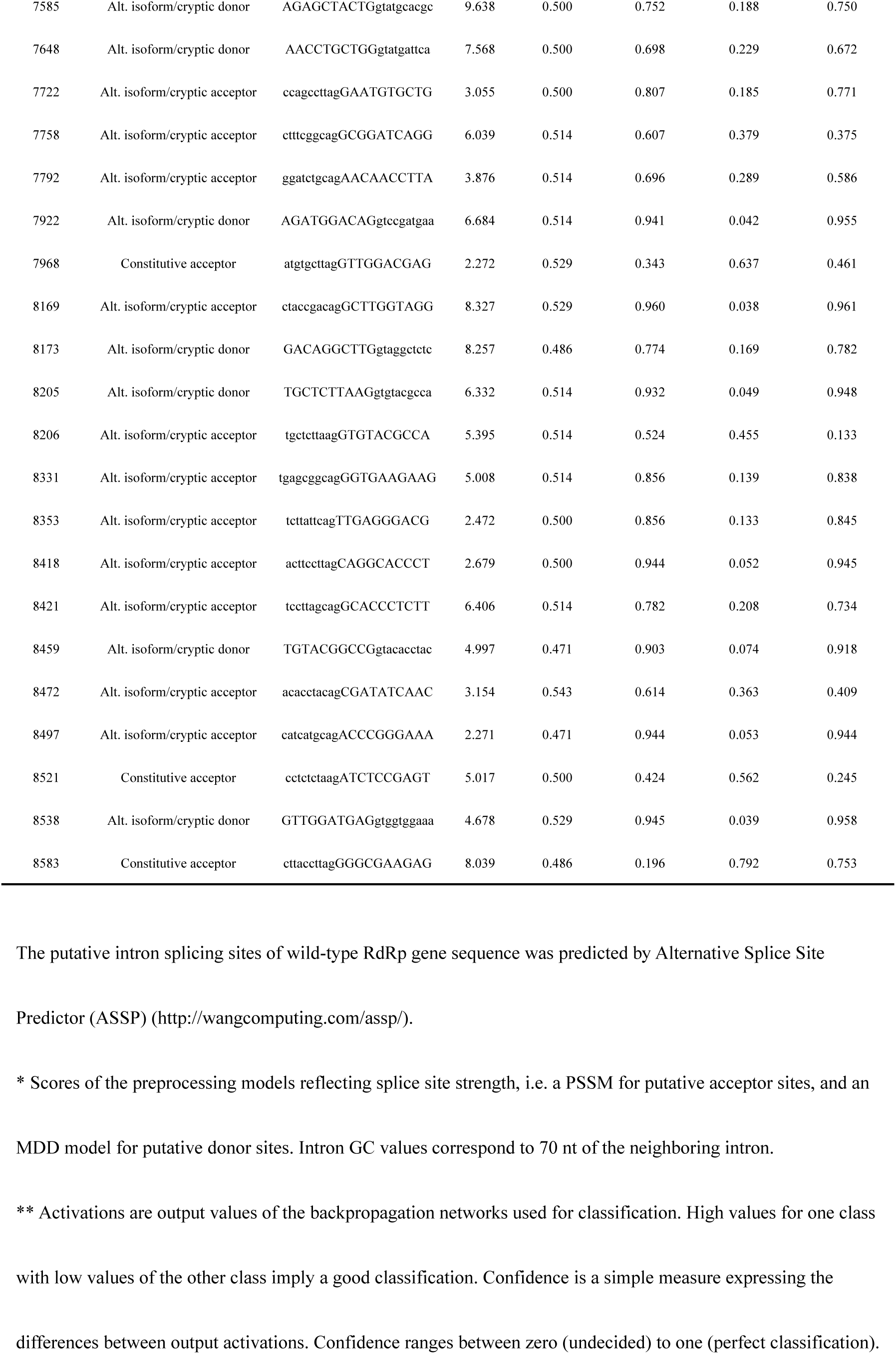
The predicted intron splicing sites of wild-type RdRp gene.

**Table S2.**
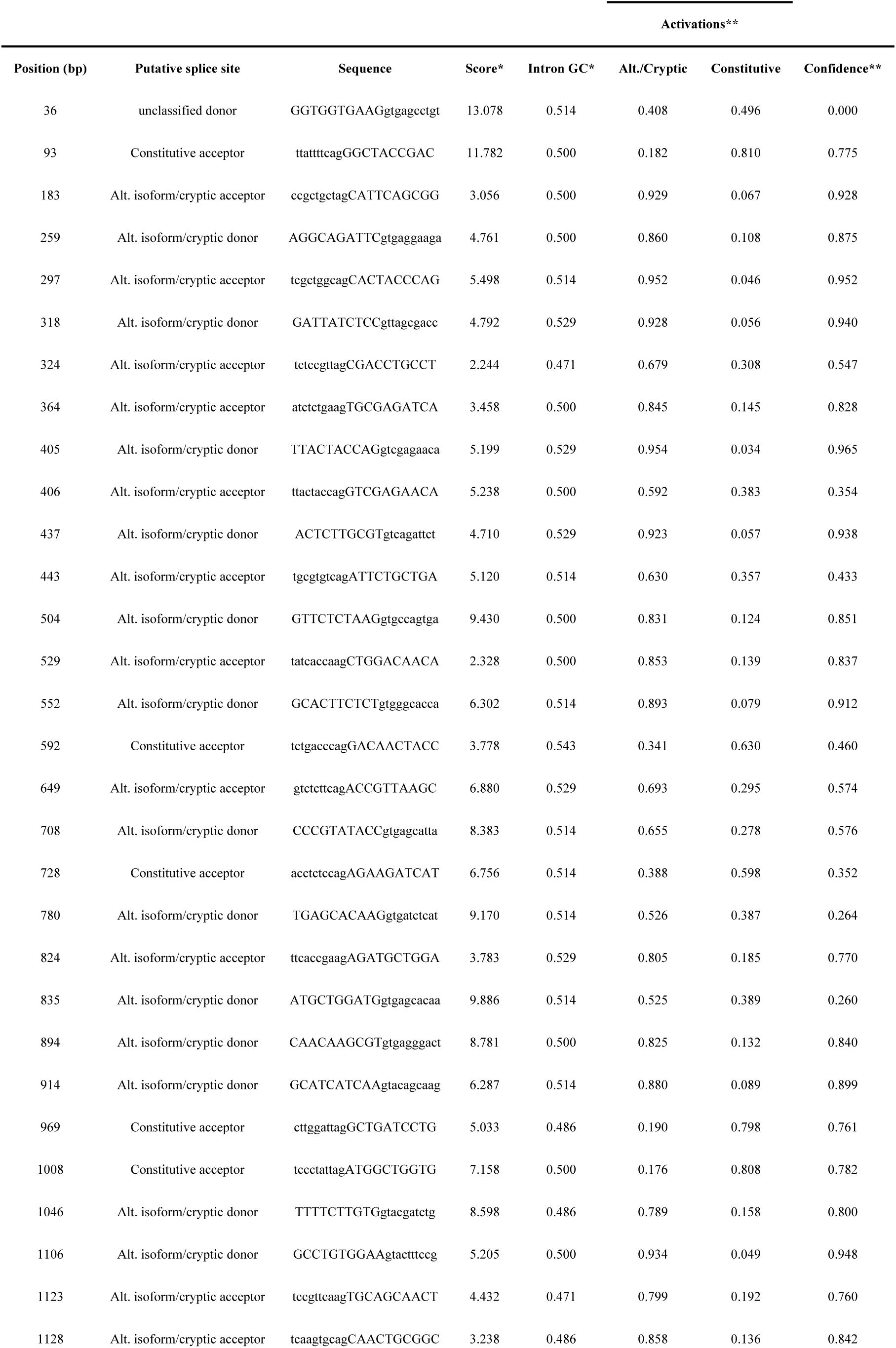
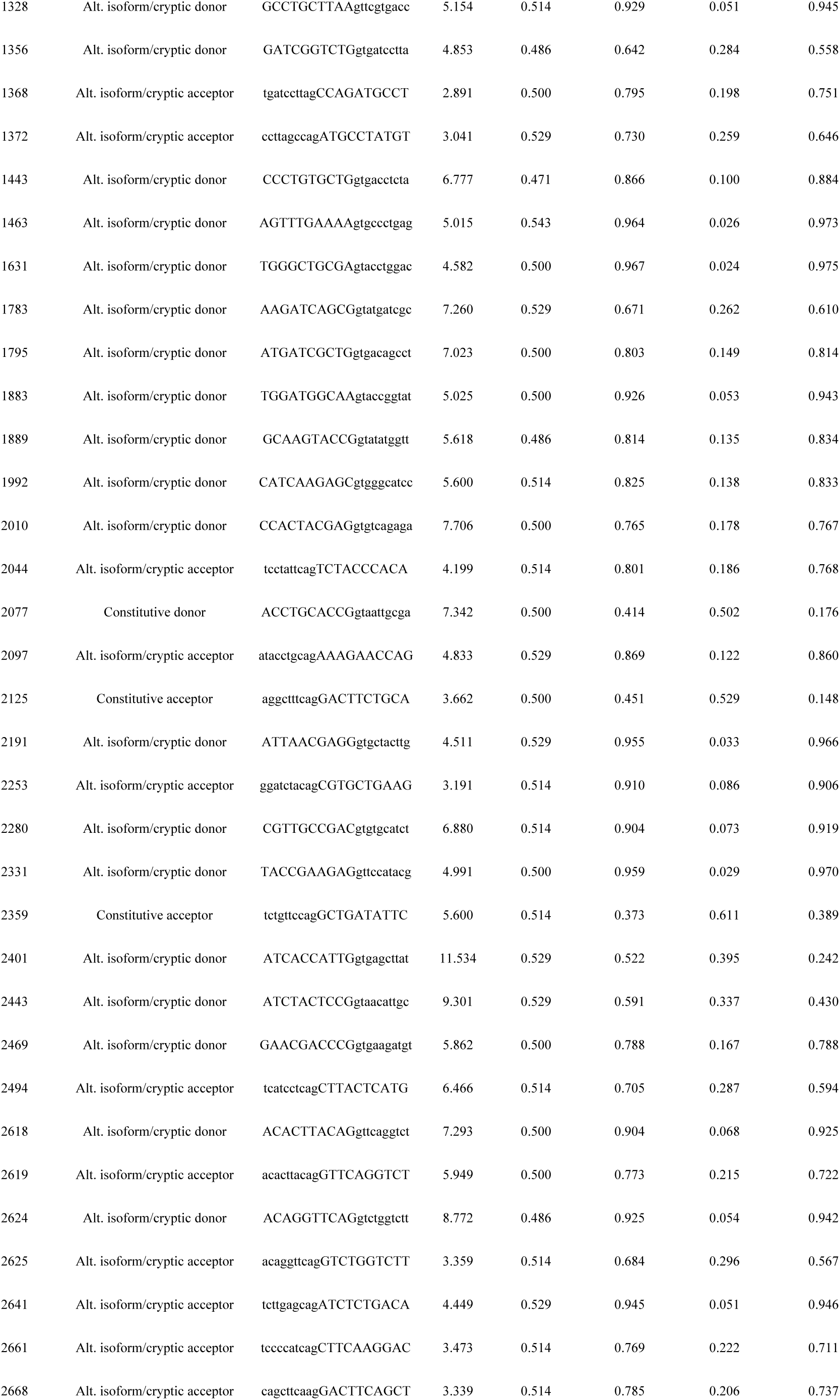
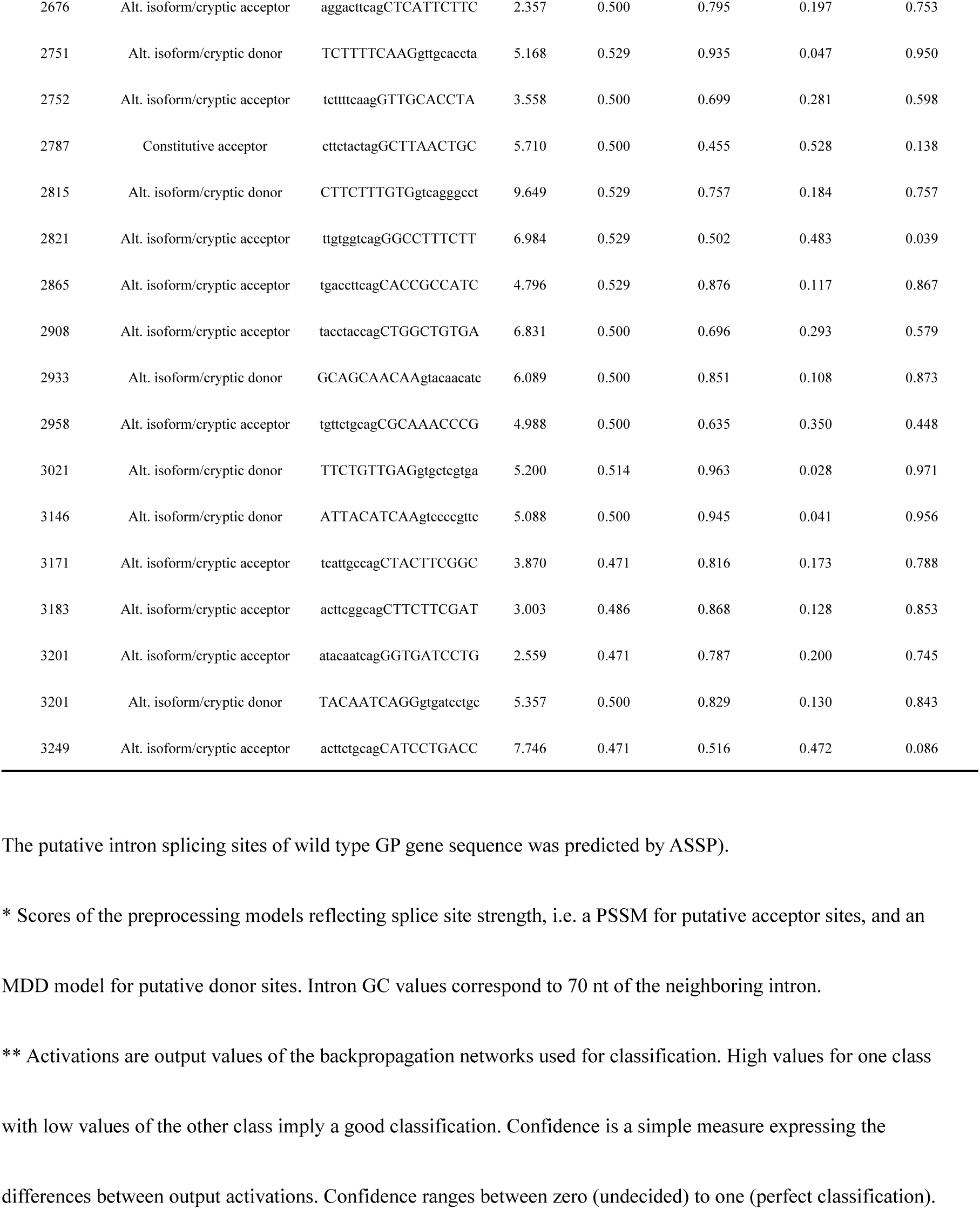
The predicted intron splicing sites of wild-type GP gene.

**Table S3.**
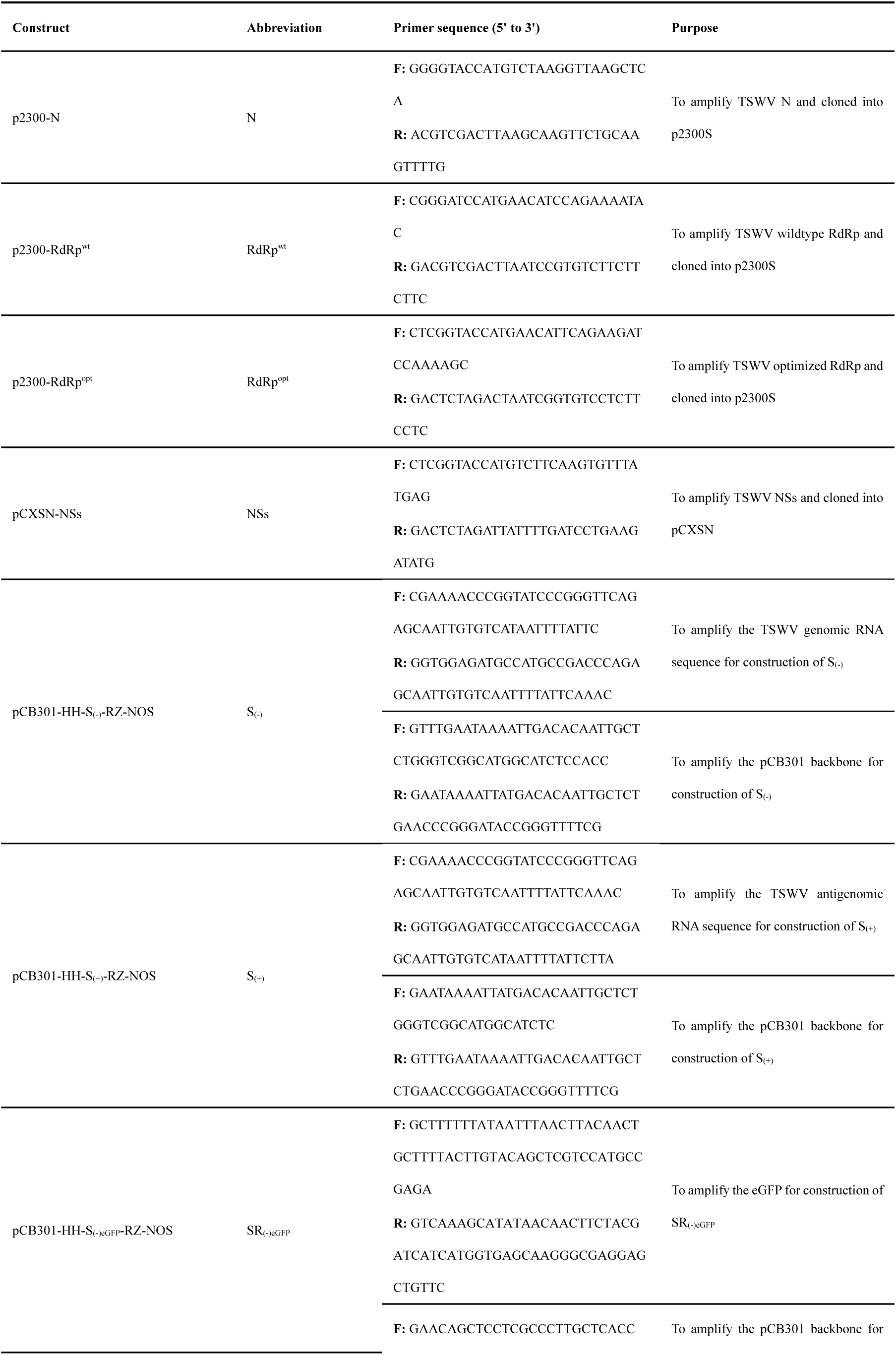
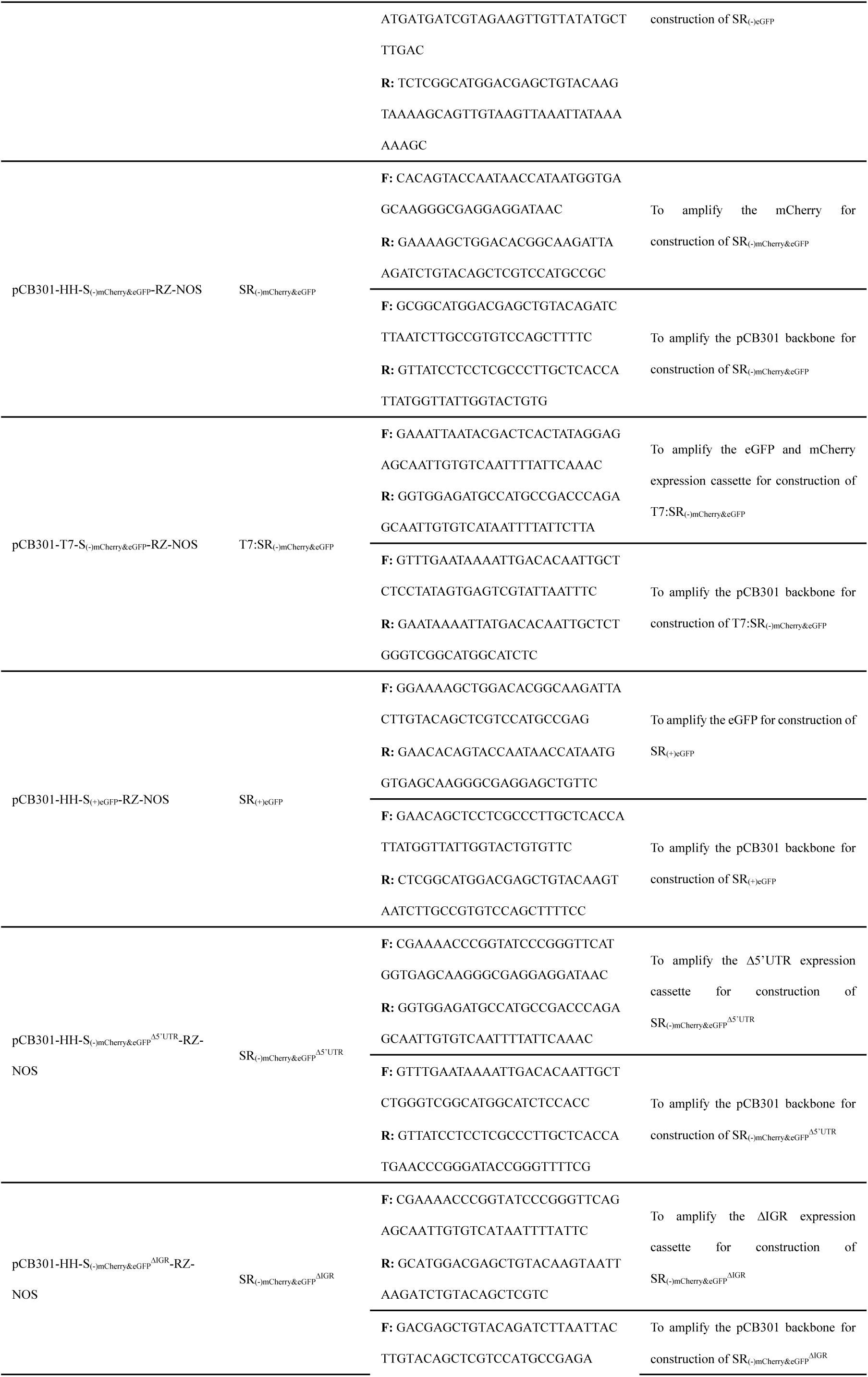
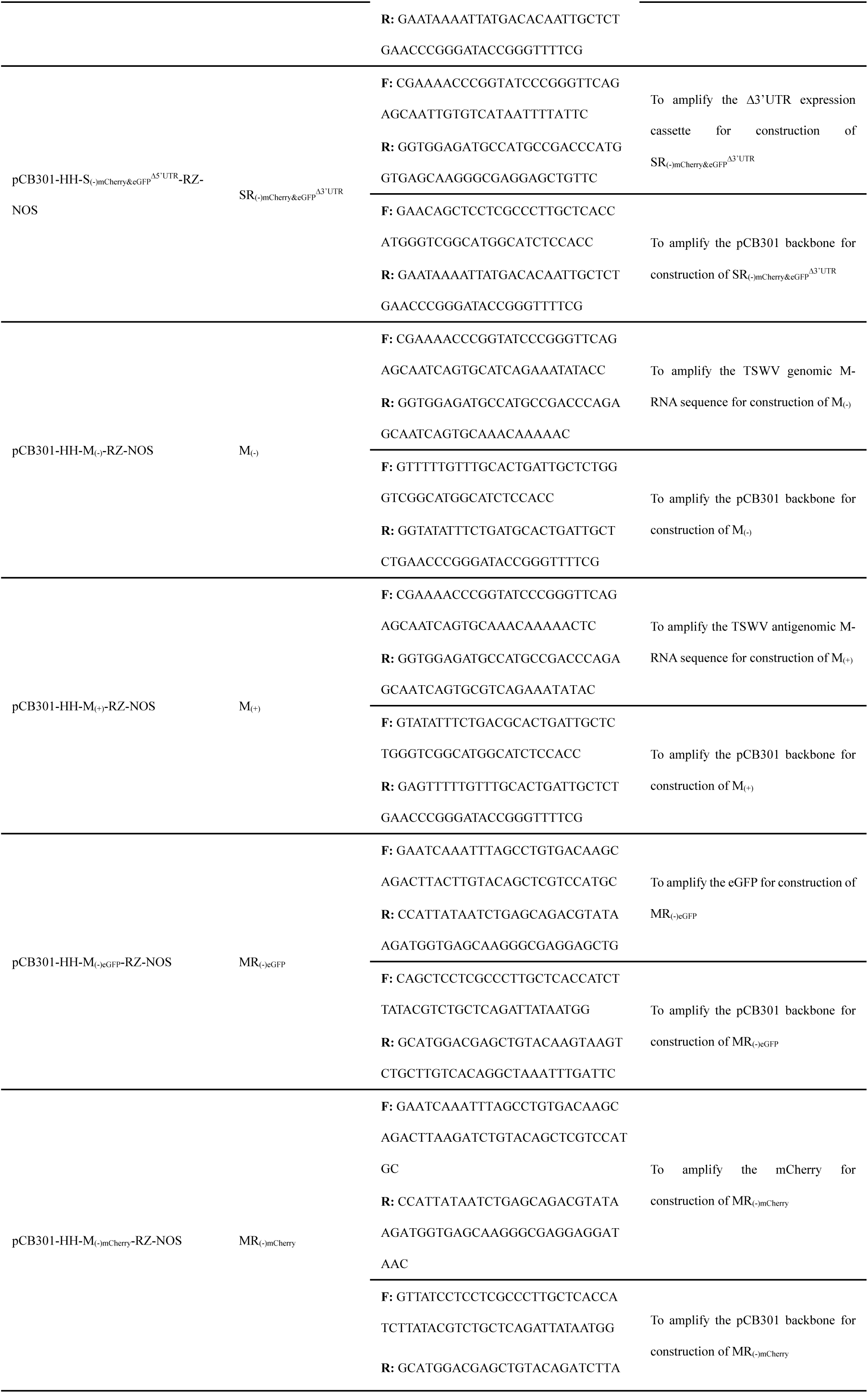
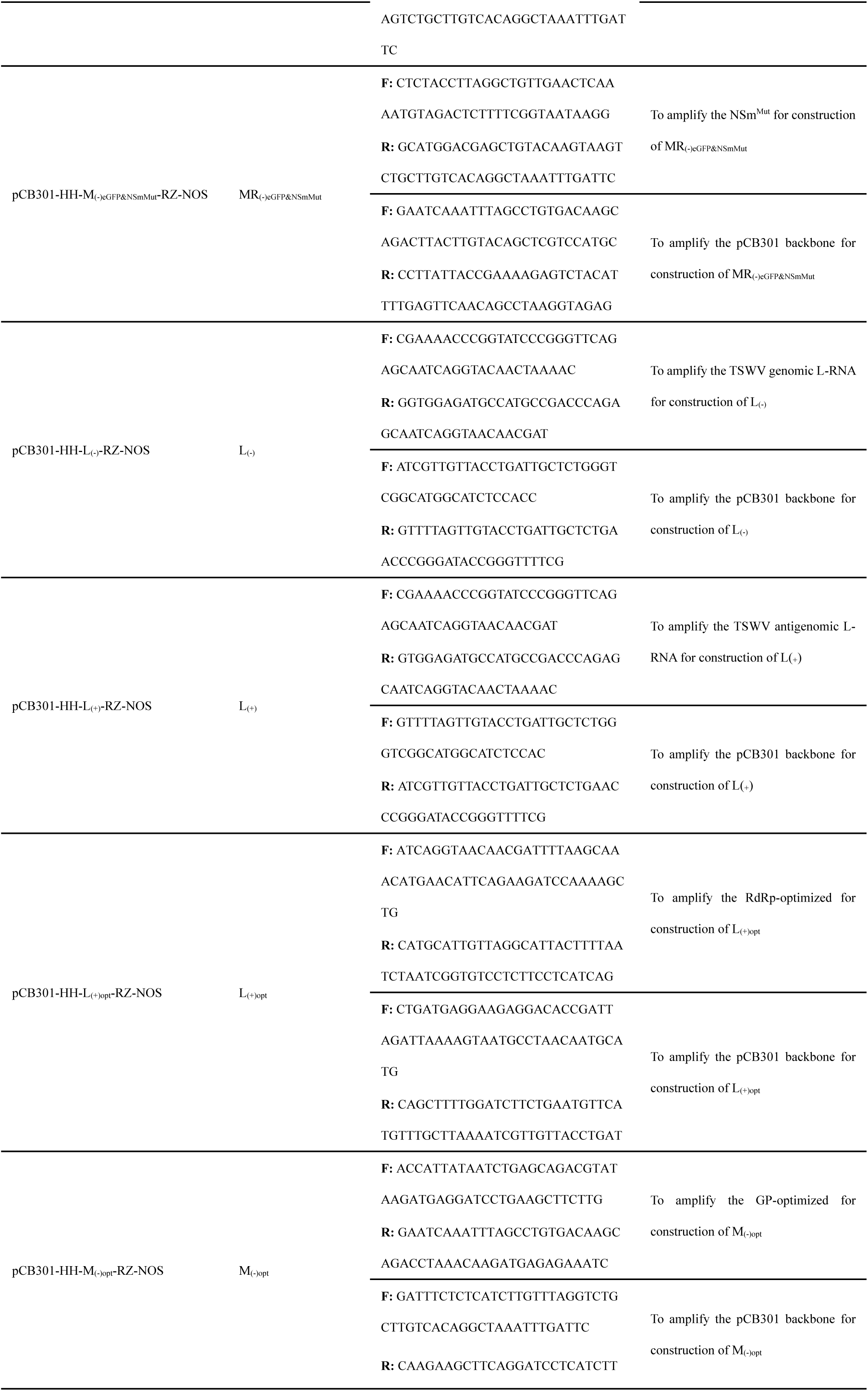
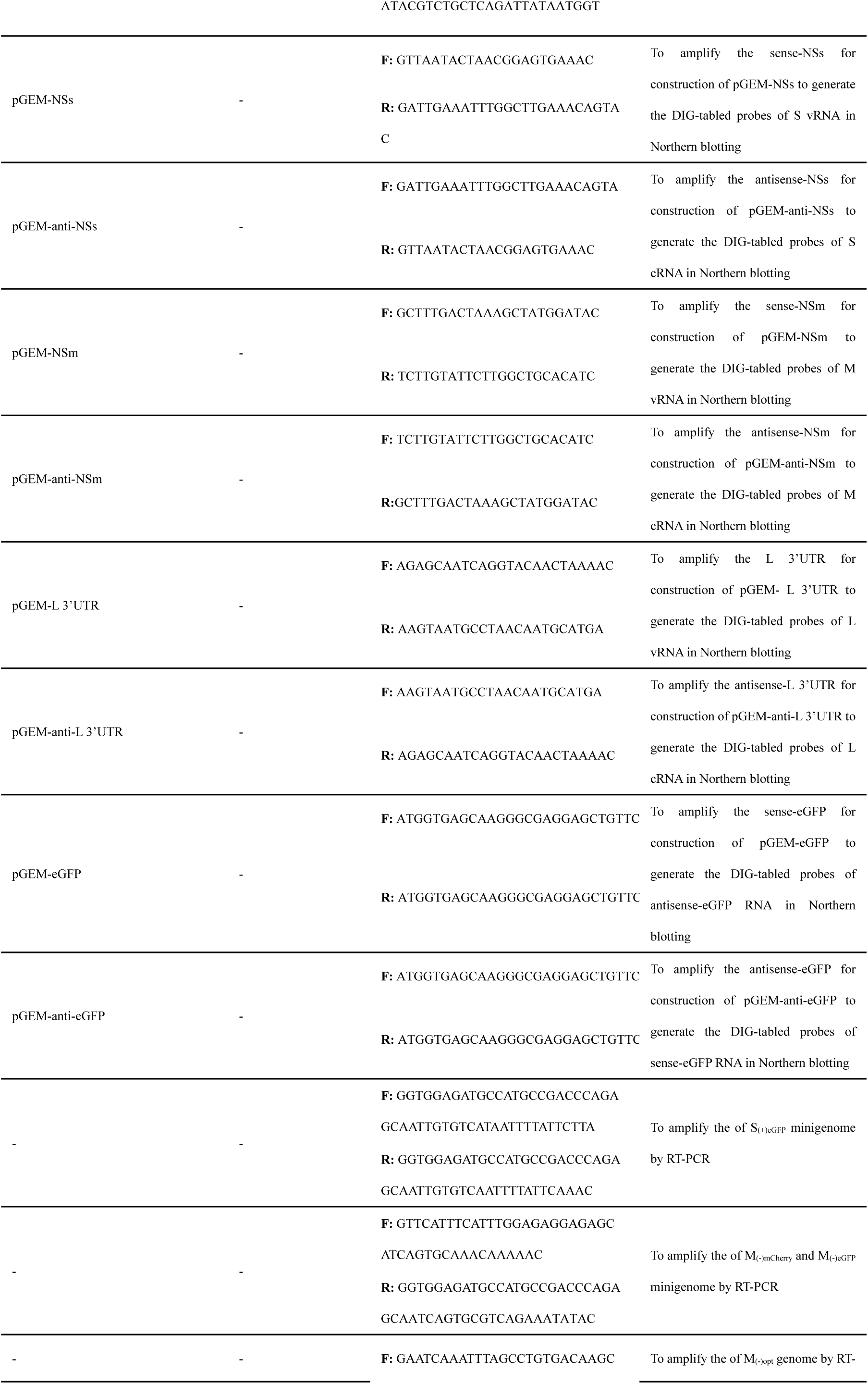
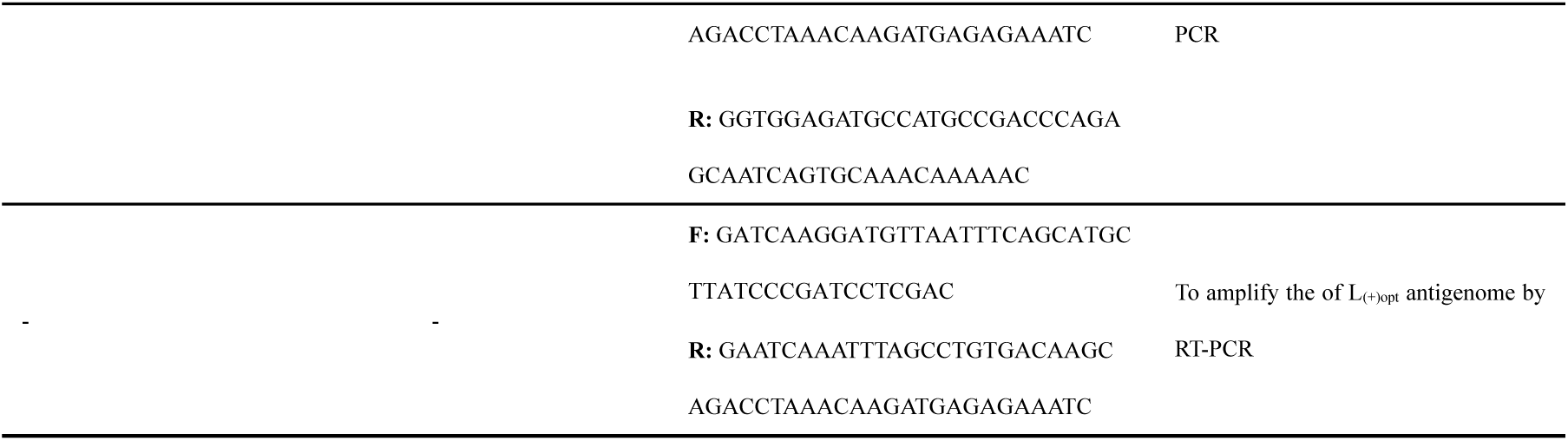
List of primers used in the study.

